# Med14 phosphorylation shapes genomic response to GLP-1 Agonist

**DOI:** 10.1101/2025.06.17.660196

**Authors:** Sam Van de Velde, Jingting Yu, K. Garrett Evensen, Edmund Pakhlevanyan, April E. Williams, Reuben J. Shaw, Marc Montminy

**Affiliations:** Peptide Biology Laboratories, The Salk Institute for Biological Studies, La Jolla CA; Razavi Newman Integrative Genomics and Bioinformatics Core, The Salk Institute for Biological Studies, La Jolla CA; Molecular and Cell Biology Laboratory, The Salk Institute for Biological Studies, La Jolla CA

## Abstract

Under feeding conditions, release of glucagon-like peptide (GLP-1) from intestinal L cells promotes insulin secretion and pancreatic beta cell viability. Binding of GLP-1 to its cognate receptor on the beta cell surface results in induction of the cAMP signaling pathway, leading to the protein kinase A (PKA) mediated phosphorylation of CREB and induction of CREB target genes. By contrast with the acute effects of this pathway on immediate early CREB target genes, which attenuate the cAMP-CREB response, sustained exposure of beta cells to the stable GLP-1 receptor agonist Exendin-4 stimulates the expression of beta cell specific CREB target genes with delayed kinetics. In a proteomic screen for transcriptional co-regulators that mediate the long-term effects of GLP-1 analogs, we identified Med14, a backbone subunit of the Mediator complex. Exposure to Exendin-4 stimulates Med14 phosphorylation at Ser983, corresponding to a conserved PKA recognition site (RRXS) that is located within an intrinsically disordered region of Med14. Phosphorylation of Med14 by PKA is essential for maintenance of enhancers that drive induction of beta cell-specific and diabetes-linked genes. Mutation of Med14 at Ser983 to alanine decreased beta cell numbers and repressed Exendin-4 induced gene regulation in primary mouse beta cells. Our work reveals how phosphorylation of a general transcription factor in response to GLP-1 analogs triggers a broad genomic response with salutary effects on beta cell function.

## Introduction

Pancreatic beta cells are equipped to convert metabolic cues of nutrient abundance to insulin secretion. Acute changes in glucose and lipid metabolism during feeding are important in maintaining energy balance; chronic increases in these circulating metabolites disrupt beta cell function, leading to beta cell failure and type 2 diabetes. Metabolic processing of glucose promotes insulin release by activating ATP synthesis followed by closure of ATP sensitive potassium (K_ATP_) channels. The attendant depolarization opens voltage gated calcium channels leading to calcium influx and release of secretory granules into the circulation [1]. Beyond its effects on insulin release, glucose promotes beta cell function by activating metabolic cycling pathways that generate secondary stimulus/secretion coupling factors and enhance excess fuel detoxification [2, 3]. Hyperglycemia stimulates glycolytic and lipogenic gene programs in beta cells through induction of Hypoxia Induced Factor (HIF) and sterol response element binding protein (SREBP) [4–9]. These responses promote adaptation to acute increases in circulating glucose concentrations, while also contributing to gluco- and lipo-toxicity in the setting of chronic hyperglycemia.

Feeding stimulates release of GLP-1 from enteroendocrine cells of the gut to maintain blood glucose homeostasis [10]. Triggering of the GLP-1 receptor in beta cells by the stable GLP-1 analog Exendin-4 (Ex-4) increases intracellular cAMP production. Ex-4 cooperates with circulating glucose levels to stimulate insulin release acutely and to increase insulin gene expression and beta cell survival [11]. The long-term metabolic benefits of Ex-4 suggest that it promotes a genomic adaptation, which enables beta cells to meet increased metabolic demands in the setting of insulin resistance [12].

Indeed, Ex-4 appears to act cooperatively with glucose to promote expression of beta cell-specific genes via the induction of CREB and its coactivators. Depletion of CREB activity in beta cells causes beta cell failure and diabetes [13]. However, the mechanisms linking gene regulation by cAMP with the salutary effects of GLP-1 analogs remain poorly understood. In a proteomic screen for transcriptional regulators that are phosphorylated by PKA in response to cAMP signaling in INS-1 beta cells, we identified MED14, a subunit of the Mediator complex.

Mediator is a multi-subunit nuclear complex that governs transcription at the level of pre-initiation complex (PIC) organization, RNA polymerase II (PolII) C-terminal domain phosphorylation, PolII pause release, and enhancer activation [14]. Structurally, mediator consists of 30 subunits that assemble into head, middle, and tail sections. The subunits are connected by the scaffolding subunit Med14. Mediator also contains a kinase module that controls activity of PolII and transcriptional elongation factors [15–18]. Aside from this kinase module, mediator lacks known enzymatic activity; rather its role in transcriptional regulation is thought to involve enhancer activation and coordination between general and DNA-specific transcription factors (TFs) through direct association with its subunits. TFs that directly bind mediator subunits include a number of metabolic regulators, such as peroxisome proliferator-activated receptor alpha and gamma (PPARα,γ) and SREBP [19–24]. Recent studies highlight a role of mediator in establishing transcriptionally hyperactive condensates containing activators, coactivators, PolII and signal-dependent TFs through liquid-liquid phase separation driven by intrinsically disordered regions (IDRs) in constituent subunits [25–27]. Enrichment of Mediator condensates over transcriptionally hyperactive genomic loci called super-enhancers appears to be essential in maintaining cell-specific gene expression and cell identity [28, 29]. These findings suggest that TF binding surfaces and IDRs in Mediator subunits could integrate metabolic signals and cell identity.

Here, we identify a conserved PKA phosphorylation site in a C-terminal IDR of Med14 that promotes sustained target gene expression by Ex-4 in INS-1 insulinoma cells through activation of CREB bound enhancers. We find that broad transcriptional regulation of metabolism coincides with cellular lipid imbalance in Med14 S983A mutant INS-1 cells. Med14 S983A mutation also blocks gene regulation by sustained Ex-4 exposure in primary beta cells. Together, our results reveal how GLP-1 analogs reprogram the beta cell genome in part through the phosphorylation of a mediator subunit.

## Results

### Exendin-4 triggers a robust transcriptional response in beta cells

To investigate the mechanism by which Ex-4 promotes insulin secretion and beta cell viability, we performed Spike-in normalized RNA sequencing (RNA-seq) on INS-1 cells upon acute (1 hour) or prolonged (16 hours) Ex-4 exposure. Ex-4 treatment for one hour induced a limited set of genes (n=184; FC > 1.5, padj. < 0.05): many of these correspond to core CREB target genes that enriched in feedback inhibition of upstream signaling (e.g. *Pde10a, Rgs2, Dusp1, Sik1*) and activator protein (AP-1) transcription factors (e.g. *Fos*, *JunB*, *Fosl1*, *Fosl2*) (**Figure 1A, Supplementary Table 1**) [30].

**Figure 1:**
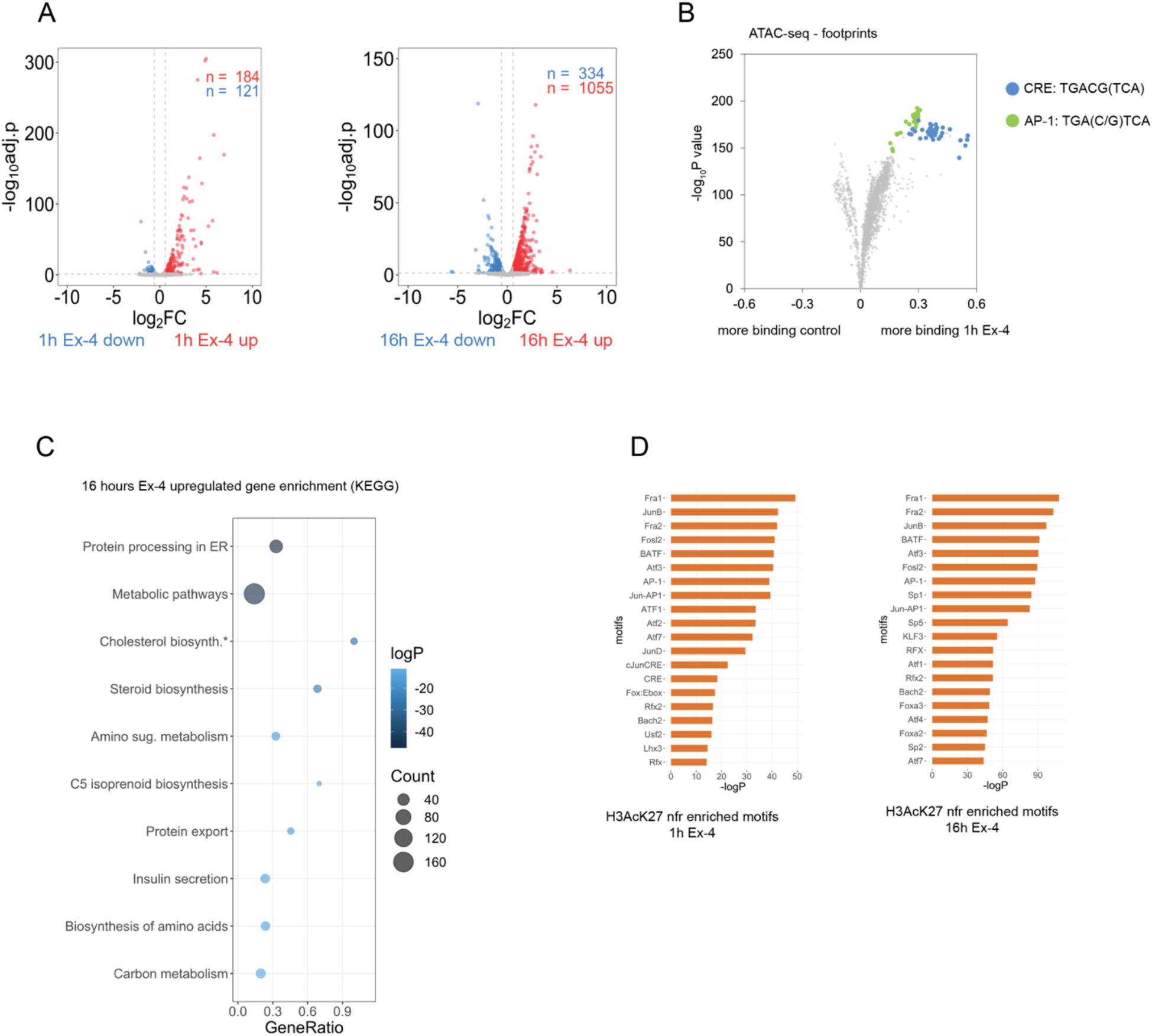
Ex-4 triggers a robust gene response. (A) Differential gene expression after acute (1h, left) and sustained (16h, right) Ex-4 (10 nM) exposure in INS-1 cells. (B) Volcano plot depicting change in footprint score over transcription factor binding motifs in accessible chromatin regions (ATAC) after 1 hour Ex-4 (10 nM) treatment. Motifs corresponding to CREB response binding (CRE) and activator protein-1 (AP-1) are highlighted. (C) KEGG pathway enrichment analysis of genes induced after 16 hour Ex-4 (10 nM) exposure. (D) Motifs enriched in nucleosome-free regions (nfr) of active enhancers (H3AcK27 ChIPseq) annotated to genes induced after 1 hour (left) and 16 hours (right).

By contrast with these short-term effects, prolonged exposure (16 hours) of INS-1 cells to Ex-4 generated a far more robust transcriptional response (n=1065; FC > 1.5, padj. < 0.05) (**Figure 1A, Supplementary Table 2,3**). Gene responses to Ex-4 overlapped almost entirely with the adenylate cyclase agonist forskolin (FSK), confirming that Ex-4 promotes gene expression primarily through activation of the cAMP pathway (**Figure S1A,B,C, Supplementary Table 4,5,6**) [31].

To determine the mechanism by which Ex-4 modulates gene expression we performed ChIP-seq studies of cAMP-responsive factors (CREB, CRTC2), activated histone marker H3AcK27, and elongating phosphorylated RNA polymerase (PolIIpS2). Inspection of the genomic region surrounding the CREB-target gene *Irs2* revealed comparable CREB/CRTC2 recruitment, enhancer activation and PolIIpS2 elongation by Ex-4 and FSK (**Figure S1D**). To further validate the role of CREB in the genomic response to Ex-4, we conducted *in vivo* footprinting analysis with high depth (∼250×10^6^ reads per sample) assay for transposase-accessible chromatin with sequencing (ATAC-seq) datasets. Footprint scoring in accessible chromatin revealed significant enrichment of CREB and CREB target AP-1 binding motifs in INS-1 cells following short-term treatment with Ex-4 and FSK (**Figure 1B, S1E,F**). Taken together, these findings confirm the importance of CREB/CRTC2 and AP-1 as drivers of gene expression by Ex-4.

The induction of core CREB target genes in response to Ex-4 is immediate and largely extinguished after 4-6 hours [32]. To investigate the role of sustained gene induction, we performed pathway analysis of transcripts induced after sustained Ex-4 and FSK exposure. Genes involved in protein processing in the Endoplasmic Reticulum (ER), metabolism, protein secretion, and cholesterol biosynthesis were up-regulated following 16h Ex-4 treatment by KEGG pathway and over-representation analysis (ORA) (**Figure 1C,S1G**). These results are consistent with a potential role for CREB in beta cell adaptation to nutrient imbalance. Motif analysis of nucleosome-free regions (NFRs) in H3AcK27 decorated active enhancers annotated to Ex-4 induced genes revealed enrichment of cAMP response elements (CREs) and AP-1 motifs after acute and sustained Ex-4 treatment (**Figure 1D**), in agreement with earlier work [33]. These findings suggest that CREB and AP-1 transcription factors play a role in both immediate and delayed gene responses to Ex-4.

### Med14 is a PKA target

We performed a proteomic screen of INS-1 cell extracts to identify regulatory proteins that are phosphorylated by PKA and that may function in modulating the transcriptional response to Ex-4. We recovered multiple Mediator subunits in Phospho-PKA substrate (RRxpS/T) antibody immunoprecipitates of INS-1 cells exposed to FSK by mass spectrometry (**Figure 2A**). Of the 10 identified Mediator components, only the scaffolding subunit Med14 was found to contain a conserved PKA substrate motif at Ser983 (in mouse) (**Figure 2B**). Ser983 is located within a 200 amino acid intrinsically disordered region (IDR) near the C-terminal tail of Med14, a region that connects the core and tail sections of mammalian Mediator (**Figure S2A**) [15, 16].

**Figure 2:**
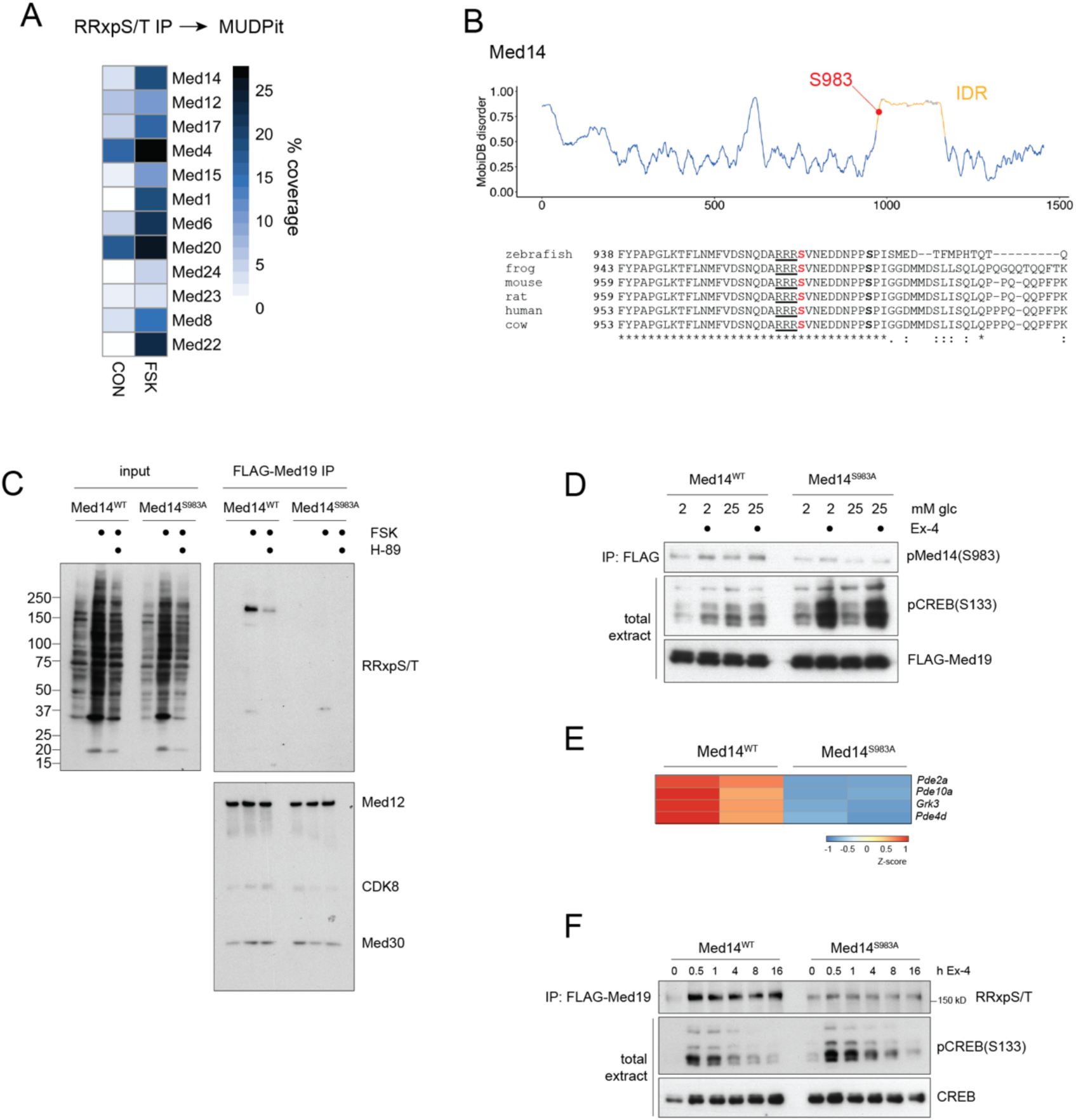
Med14 is a PKA target. (A) MUDPit coverage of mediator complex subunits recovered in PKA substrate (RRxpS/T) immunoprecipitates from control and FSK (10 μM) treated cell lysates. (B) MobiDB disorder score over Med14. S983 is indicated inside the IDR (top). Alignment of Med14 orthologs highlighting conservation of the PKA substrate motif and the Erk target site Ser992 (ref) (bottom). (C) Western blot of PKA substrate antibody on input (left) and endogenous FLAG-Med19 immunoprecipitates (IP) (right) in Med14^WT^ and Med14 S983A Crispr knock-in lines. Integrity of mediator complex in IP is probed by Med12, CDK8 and Med30 blots. PKA activation by FSK and inhibition by H-89 is shown in total extract. (D) Med14S983 and CREBS133 phosphorylation by Ex-4 (10 nM) in WT and Med14 S983A mutant lines. FLAG-Med19 is probed as loading control in total extract. (E) Expression of cAMP signal-attenuating genes is repressed in Med14 S983A mutant. (F) Time-course western after treatment of WT and Med14 S983A cells with Ex-4 (10 nM). FLAG-Med19 immunoprecipitates are probed with PKA substrate (RRxpS/T) antibody. Phosphorylated (pS133) and total CREB are probed in total extract.

To address whether Med14 S983 is phosphorylated by PKA, we generated independent Med14 Ser983Ala mutant INS-1 lines (see methods). We also generated endogenous FLAG-Strep-Med19 fusions in WT and Med14 S983A INS-1 backgrounds to isolate WT and mutant Mediator complexes. FLAG-Med19 immunoprecipitates from WT and Med14S983A extracts followed by immunoblot with phospho-PKA substrate antibody confirmed phosphorylation around the predicted molecular weight of Med14 (∼160kD) in WT, but not mutant Med14 cells. Moreover, Med14 phosphorylation was blocked by the PKA inhibitor H-89 (**Figure 2C**).

In principle, Med14 S983 mutation could destabilize Mediator and disrupt interaction of Med19 with other Mediator subunits, thereby impeding detection of phospho-PKA substrate immunoreactivity in other Mediator constituents or interacting TFs. Arguing against this notion, immunoblot for additional Mediator subunits (Med12, Cdk8 and Med30) in FLAG-Med19 immunoprecipitates revealed comparable amounts of these proteins in WT and Med14 S983A mutant complexes (**Figure 2C**). We further confirmed the integrity of Med14 S983A mutant Mediator by glycerol gradient sedimentation of FLAG-Med19 immunoprecipitates (**Figure S2B**). Taken together, these findings show that Med14 is the predominant PKA substrate in the Mediator complex.

To determine whether Ex-4 modulates Med14 phosphorylation, we generated a phospho-specific antibody targeting the Med14pS983 site. In the basal state, WT and S983A mutant Med14 were undetectable by immunoblot using the phospho-specific antiserum. Ex-4 treatment increased amounts of phospho-Med14 in WT but not mutant Med14 in both low (5mM) and high (20mM) glucose media. By contrast, exposure of Med14 S983A cells to Ex-4 strongly induced CREB hyperphosphorylation in response to Ex-4, suggesting that Med14, like CREB, plays a critical role in repression of upstream cAMP signaling (**Figure 2D**). Among the genes repressed in Med14S 983A lines, we identified phosphodiesterases (*Pde2a*, *Pde10a*, *Pde4d*), as well as the G-protein coupled receptor kinase *Grk3*, which was recently identified as a negative regulator of the GLP-1 receptor (**Figure 2E**) [34, 35]. Consistent with the notion that CREB activates immediate-early gene expression, stimulus-induced CREB phosphorylation decreases upon prolonged stimulation through dephosphorylation by PP1 and PP2A [36, 37]. By contrast, phosphorylation of Med14 by Ex-4 was sustained, even after 16 hour stimulation (**Figure 2F**). Together, these results suggest that phosphorylation of CREB and Med14 cooperatively promote gene induction by Ex-4 on different timescales.

### Med14 S983 phosphorylation promotes target gene induction through activation of relevant enhancers

We next explored the role of Med14 phosphorylation in promoting Ex-4-induced gene expression. RT-QPCR analysis of core CREB target genes in WT and Med14 S983A mutant INS-1 lines revealed that *Fos* induction peaked at both protein and mRNA levels after 1 hour and returned to baseline within 4-8 hours at both mRNA and protein levels (**Figure 3A,3B**). Further, Ex-4- and FSK-induced activity of a CRE-luciferase reporter was blunted when expressed in Med14S983A mutant cells (**Figure S3A**). Notably, delayed-early gene (DEG) induction (*Kdr*, *Osbpl6*) was quantitatively blocked in Med14 S983A mutant lines exposed to Ex-4 (**Figure 3A**). To confirm the role of cAMP in mediating these effects, we repeated time-course induction of DEGs using FSK. *Kdr*, *Osbpl6* and tissue-type plasminogen activator (*Plat*) genes were substantially up-regulated following activation of the cAMP-PKA pathway in INS-1 cells (**Figure S3B**). In contrast, triggering of the NFκB pathway in response to IL-1β actually increased in Med14 S983A mutant cells relative to WT, supporting the notion that Med14 phosphorylation has selective effects on cAMP inducible and pro-inflammatory genes (**Figure S3C**).

**Figure 3:**
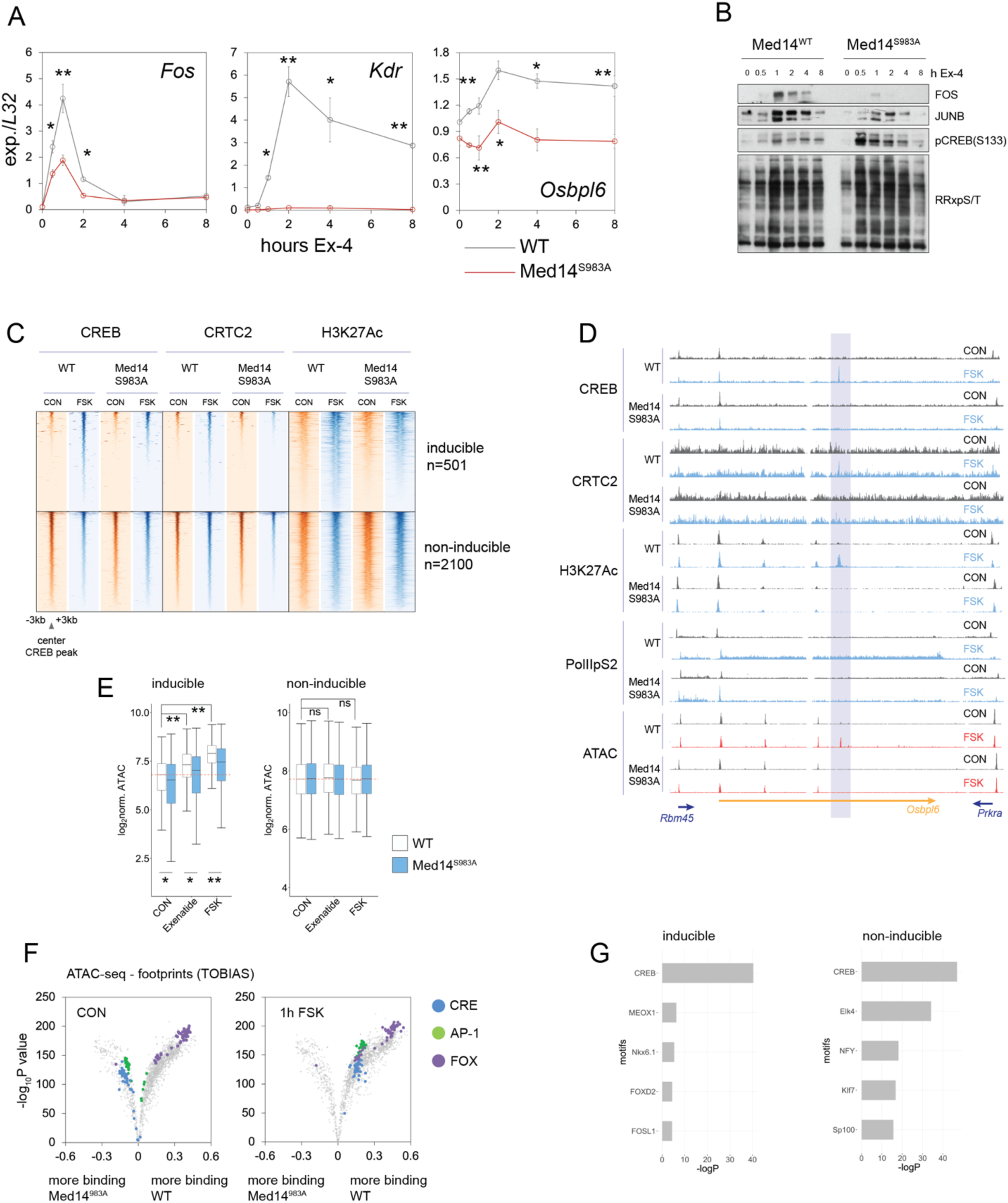
Med14S983 phosphorylation promotes gene induction by activating enhancers. (A) Time course Q-PCR for immediate-early (*Fos*) and delayed-early (*Kdr*, *Osbpl6*) genes over 8 hour Ex-4 (10 nM) treatment in WT and Med14 S983A mutant cells. Error bars show standard deviation. *p<0.05, **p<0.01 by unpaired t test. (B) Time course western blot for core CREB target gene products FOS and JUNB, phospho-CREB(S133) and phospho-PKA target over 8 hour Ex-4 (10 nM) treatment. (C) Chromatin occupancy (ChIP-seq) of CREB, CRTC2 and H3AcK27 in WT and Med14 S983A mutants in basal and FSK-stimulated conditions over cAMP-inducible (top) and non-inducible (bottom) enhancers. (D) Example of an inducible enhancer in the *Osbpl6* intron. Tracks are scaled across conditions for each ChIP target and for ATAC. Intronic cAMP responsive enhancer is highlighted. (E) Box plots comparing ATAC-seq read counts over cAMP-inducible (left) and non-inducible (right) enhancer regions. (F) Volcano plot comparing change in binding over known transcription factor binding motifs between WT and Med14 S983A mutants in control (left) and FSK (10 μM) treated (right) cells. Footprinting analysis was performed on high-depth (∼250×10^6^ reads per sample) ATAC-seq data. (G) Enrichment of transcription factor binding motifs over cAMP-inducible (left) and non-inducible (right) enhancers.

cAMP-inducible gene expression is dependent on CREB occupancy over stimulus-responsive promoter-distal enhancers, which are defined by co-recruitment of CBP/p300 and subsequent activating acetylation of histones H3 and H2A.Z [38, 39]. Of the 4200 CREB bound enhancer regions in INS-1 cells, only ∼12% (501) exhibited increases in H3AcK27 following exposure to FSK, suggesting that additional nuclear factors are required for stimulus-induced transcription (**Supplementary Table 7,8**). Mediator has been found to promote transcription from tissue-specific rather than global enhancers. We considered that, following their phosphorylation by PKA, CREB and Med14 may act cooperatively on Mediator sensitive enhancers. Inducible enhancer activity at CREB bound genomic loci was significantly reduced in MED14 S983A mutant cells relative to WT. ChIPseq data for H3AcK27 revealed a reduction in inducible enhancer activity in MED14 S983A mutant INS-1 cells, while enhancer activity was unchanged either globally or across non-inducible CREB bound enhancers. In line with this result, CREB and CRTC2 occupancy in Med14 S983A mutant cells are decreased over inducible enhancers in response to FSK (**Figure 3C,D,S3D**).

Tissue-specific enhancers exhibit enhanced nucleosome clearance compared to ubiquitous regulatory regions in the genome [40]. We examined the contribution of Med14 phosphorylation on DNA accessibility in enhancers by ATAC-seq. As expected, 1 hour Ex-4 or FSK treatment increased chromatin accessibility over cAMP responsive, but not non-inducible enhancers (**Figure 3D,3E,S3E**). Stimulus-induced accessibility was blunted in Med14 S983A mutant cells. Genome-wide footprinting analysis on ATAC-seq data revealed significant enrichment of CREB binding motifs after acute Ex-4 or FSK stimulation, consistent with a role for CREB in mediating the immediate early gene response to cAMP (**Figure 3F, S3F**). Surprisingly, binding over CRE and AP-1 motifs was enriched in Med14 S983A relative to WT cells under basal conditions by footprinting analysis. In line with this observation, FOXO motifs were also enriched in inducible, but not non-inducible CREB bound loci (**Figure 3G**).

### Med14 S983 is a critical mediator of insulin gene expression

Given the apparent role of mediator for induction of cell-specific genes, we compared expression of beta cell-restricted RNAs (MSigDB Hallmark pancreatic beta cells n=37) in WT and Med14 S983A cells. Med14 mutation resulted in a small but significant decrease in beta cell-specific transcripts that was more pronounced after 16 hours Ex-4 exposure (**Figure 4A**).

**Figure 4:**
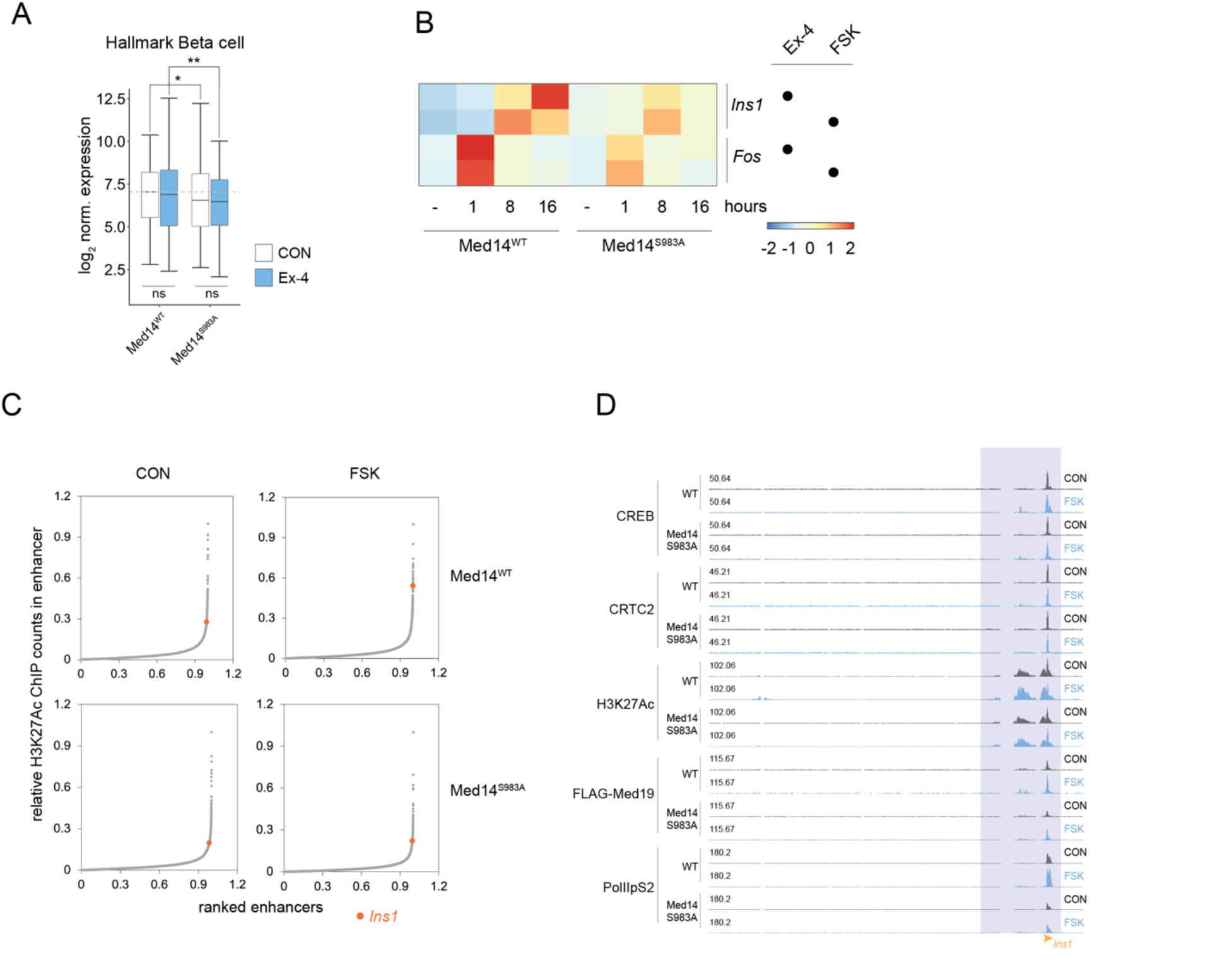
Med14S983 is a critical mediator of insulin gene induction. (A) Normalized expression of beta cell-specific genes (MSIGDB Hallmark) between WT and Med14 S983A mutant cells. (B) Induction profile of *Ins1* and *Fos* by Ex-4 (10 nM) or FSK (10 μM) in WT and Med14 S983A mutants. (C) Ranked enhancer score (H3AcK27 ChIPseq) plots of WT (top) and Med14 S983A mutant cells (bottom) in basal or FSK stimulated state. Insulin (*Ins1*) is highlighted. (D) ChIPseq tracks at the *Ins1* locus in basal and FSK stimulated states comparing ChIP occupancy between WT and Med14 S983A mutant cells.

Insulin is the most highly expressed cell-specific gene in beta cells and its induction by cAMP is thought to promote beta cell adaptation to increases in circulating glucose and fatty acids. By contrast with the rapid induction of the Fos gene (1 hour) by Ex-4 and FSK, *Ins1* gene expression peaked after 16 hours in response to both stimuli; these effects were blunted in Med14 S983A mutant cells (**Figure 4B**). Indeed, ranked enhancer strength plots revealed induction of *Ins1* enhancer activity upon FSK exposure in WT but not Med14 S983A mutant cells, indicating that Med14 S983 phosphorylation may be required for stimulus-induced *Ins1* expression through enhancer activation (**Figure 4C**). Further inspection of the *Ins1* locus showed reduced activation of two proximal enhancers, as well as decreased recruitment of CREB, CRTC2, and Mediator over proximal genomic regions (**Figure 4D**).

### Ex-4 tunes metabolism through Med14

To further explore the role of Med14 phosphorylation in Ex-4-driven beta cell transcription, we compared RNA-seq datasets of WT and Med14 S983A INS-1 cells treated with Ex-4 for 1, 8 or 16 hours. The transcriptional response to sustained (16 hours) Ex-4 exposure was significantly blunted in Med14 S983A mutant cells (**Figure 5A, Supplementary Table 9,10,11**); but global transcription was relatively unaffected in Med14 S983A mutants (**Figure S5A**), suggesting that MED14 acts on a subset of genes that function in ER stress, metabolism, and insulin secretion (**Figure 5B**). Cluster analysis of genes uniquely induced in WT cells after 16h Ex-4 treatment (n=636; FC > 1.5, padj. < 0.05) revealed that S983A mutation of Med14 reduces the response to Ex-4 in part by increasing the basal expression of these genes (**Figure S5B, clusters 1 and 2**). Consistent with this finding, KEGG analysis revealed significant overlap in pathway enrichment between genes induced upon prolonged Ex-4 exposure and genes up-regulated in Med14 S983A mutants under basal conditions (**Figure 1C, S5C,D**).

**Figure 5:**
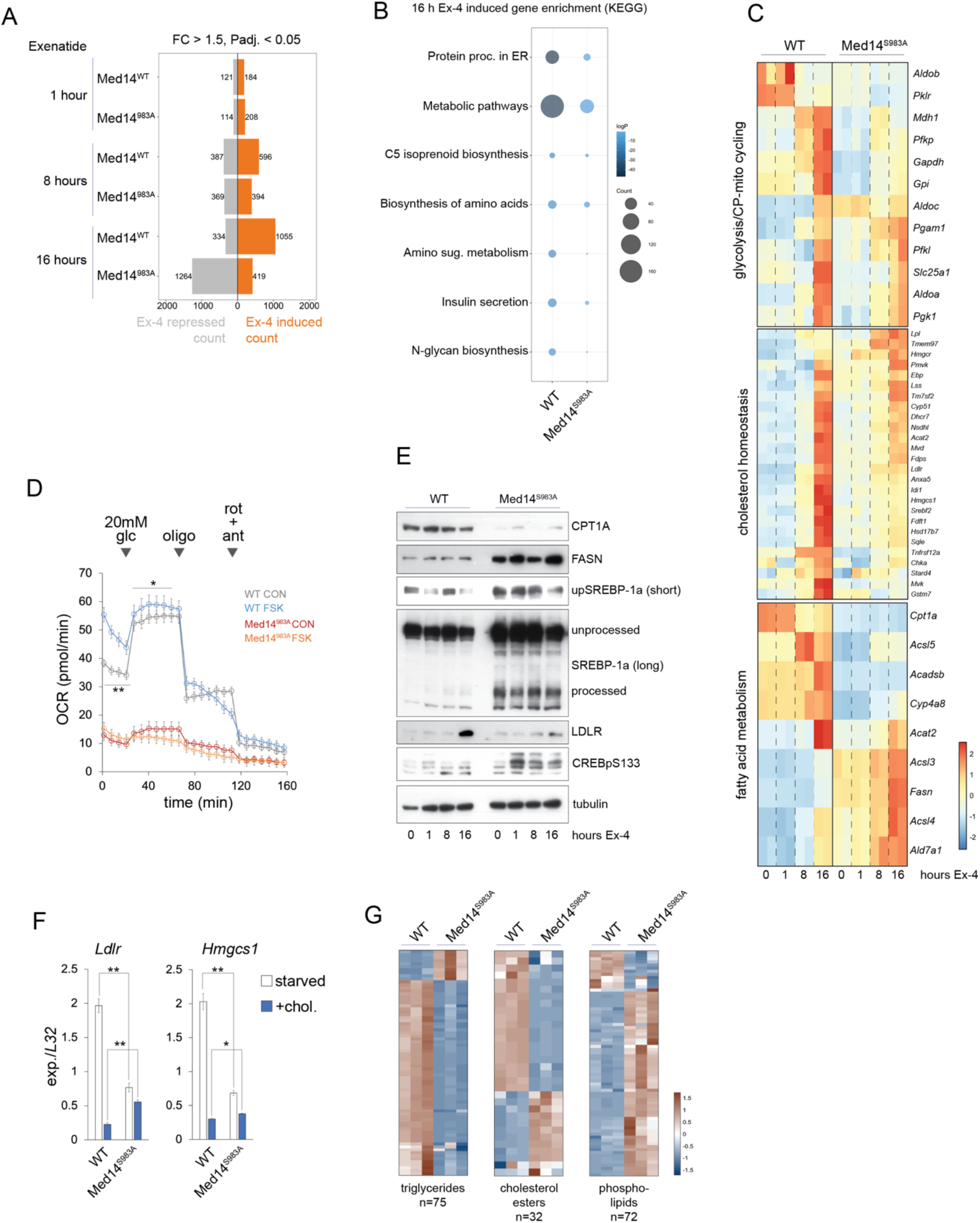
Ex-4 tunes metabolism through Med14. (A) Ex-4 regulated gene counts from spike-in corrected RNA-seq after 1, 8 and 16 hour exposure in WT and Med14 S983A mutant cells. (B) KEGG pathway enrichment of Ex-4-induced genes (16 hours) in WT and Med14 S983A mutant cells. (C) Heatmaps of metabolic gene regulation by Ex-4. Glycolytic genes with *Mdh1* and *Slc25a1* (top), cholesterol homeostasis genes (middle) and fatty acid metabolism genes (bottom). (D) Oxygen consumption rate (OCR) of WT and Med14 S983A mutants with or without 16 hour pre-exposure to FSK (10 uM). Basal glucose concentration is 2 mM. Cells were treated with 20 mM glucose, 5 uM oligomycin and 5 uM rotenone + 5 uM antimycin A at the indicated timepoints. (E) Western blots probing fat metabolism and CREB activation in WT and Med14 S983A mutant cells. Cells were exposed to Ex-4 as indicated. Unprocessed and cleaved SREBP-1a bands are marked. (F) RT-qPCR of *Ldlr* and *Hmgcs1* comparing WT and Med14 S983A mutant cells. Cells were cultured for 4 days in lipoprotein deficient serum and treated with 10 ug/ml cholesterol plus 1 ug/ml 25-hydroxycholesterol for 8 hours as indicated. * p < 0.002, ** p < 0.0003 by unpaired t test. (G) Heatmap comparing levels of triglycerides (left), cholesterol esters (middle) and phospholipids (right) in WT and Med14 S983A cells.

Beta cells respond to increases in circulating glucose concentrations by inducing glycolytic and lipogenic gene expression. However, chronic overexpression of glycolytic and lipogenic programs is also associated with disruption of metabolism-secretion coupling, beta cell failure, and diabetes, indicating that appropriate regulation of metabolic genes is crucial for beta cell function. Having seen enrichment of metabolic genes after sustained Ex-4 treatment in standard RPMI media (11mM glucose), we next evaluated the role of Med14 S983 in regulation of glycolytic gene expression. Glycolytic gene induction by Ex-4 is mostly suppressed in Med14 S983A mutant cells. In addition, genes involved in cyclic pyruvate pathways, cytoplasmic malate dehydrogenase 1 (*Mdh1*) and the mitochondrial citrate carrier (*Slc25a1*), are strongly induced in WT, but not mutant cells. (**Figure 5C**). Moreover, pyruvate dehydrogenase kinase genes *Pdk1* and *Pdk2* are down-regulated in mutant cells, indicating increased entry of pyruvate into the TCA cycle and several TCA cycle genes are induced as well (*Sucla2*, *Idh*, *Sdhc*, *Ogdh*) (**Figure S5E**). These observations suggest that Ex-4 can accelerate metabolic pathways that are linked to insulin secretion. To test this idea, we measured oxygen consumption rate (OCR) in WT and mutant cells exposed to FSK for 16 hours. Treating WT cells with FSK induced OCR under low (5 mM) and high (20mM) glucose conditions. In contrast, OCR was substantially reduced in Med14 S983A cells and no significant response to FSK or high glucose concentrations (**Figure 5D**). Together, these results suggest that Med14 phosphorylation supports accelerated glucose metabolism and ATP-linked respiration upon Ex-4 exposure.

Lipogenic and cholesterol synthetic genes were induced in WT INS-1 cells after prolonged EX-4 treatment in WT cells (**Figure 5C middle and bottom**). Basal expression of these genes was mostly increased in Med14 S983A mutants, while activation after 16 hours Ex-4 was blunted, indicating that Med14 S983A modulates lipogenic genes in both basal and stimulated states (**Figure 5C middle and bottom**). Western blot analysis revealed that Ex-4 treatment resulted in increased SREBP-1 cleavage in WT but not Med14 S983A mutant cells. In keeping with increased basal cholesterol synthesis gene expression, full-length and processed SREBP-1 were substantially increased in Med14 S983A mutant cells versus WT (**Figure 5E**). However, these increases in activated SREBP-1 levels were not sufficient for induction of SREBP target gene protein levels in mutant cells, consistent with earlier work showing that differences in metabolic gene expression are often more pronounced at the protein rather than the transcript level (*Ldlr*; **Figure 5E**) [41]. Nutrient-induced SREBP target gene expression was disrupted in Med14 S983A mutant cells as well. Indeed, expression of *Ldlr* and *Hmgcs1* was significantly lower in lipid starved Med14 S983A mutant cells compared to WT, while repression by cholesterol treatment was blunted, resulting in higher SREBP target gene expression in non-starved conditions. Together, these data indicate Med14 S983 plays an important role in promoting lipid homeostasis in part by enabling SREBP target gene regulation.

We noticed that increased cleavage of SREBP −1 in mutant cells coincided with induction of acyl-CoA synthases (*Acss*, *Acsl*) and *Fasn,* and suppression of genes involved in fatty acid and branched chain amino degradation (*Cpt1a, Acadsb, Cyp4a8*) (**Figure 5C, bottom, 5E**). These results suggest an imbalance in intracellular energy stores in mutant cells. To test this possibility, we compared lipid profiles of WT and Med14 S983A mutant cells by untargeted lipidomics screening. Surprisingly, storage lipids (triglycerides, free fatty acids and cholesterol esters) were mostly depleted in mutant cells, while phospholipids and diacylglycerol were generally increased (**Figure 5G, Supplementary Table 12**).

### Med14S983 controls gene activation by Ex-4 in primary mouse islets

To determine the role of Med14 S983 in pancreatic islets, we undertook single nucleus RNA-seq (snRNA-seq) of cultured islets isolated from WT and whole-body knock-in Med14 S983A mutants. We identified alpha, beta, delta and PP cell populations in WT and mutant tissue (**Figure 6A**). Relative beta cell numbers were significantly reduced while alpha and delta cell content increased in Med14S983A mutant islets, similar to the shift in alpha-to-beta cell ratio observed in Type 2 diabetes (**Figure 6B**)[42, 43]. Pseudobulk analysis of gene expression in beta cells revealed repression of several cell type-specific genes in mutant beta cells (e.g. *Iapp*, *Slc2a2*, *Slc30a8*), while glucagon (*Gcg*) gene expression was up-regulated in mutant beta cells (**Figure 6C, Supplementary Table 13**). KEGG pathway analysis revealed repression of pathways involved in growth, differentiation, secretion, and cell adhesion (*mTOR, MAPK, Rap1, Wnt, EGFR*) in Med14 S983A mutants, suggesting that impaired expression of signaling molecules underlies diminished growth and differentiation in mutant beta cells (**Figure 6D**). Consistent with our observations in INS-1 cells, negative modulators of cAMP signaling (*Grk5, Pde3b*) were repressed in Med14S983A mutant islet cells, confirming that Med14 is an effector of the cAMP pathway in primary cells (**Figure S6A**). Furthermore, mediator subunits *Cdk8*, *Med12* and *Med25*, activating histone H3 methyl transferases (*Kmt2a, Ehmt2, Setd1a, Dot1l*) as well as genes controlling posttranscriptional processing of mRNA (splicing, mRNA transport and surveillance, exon junction complex) are induced in Med14 S983A mutant cells, while the conserved non-coding RNA 7SK, a component of the repressor to the transcription elongation complex P-TEFb, is repressed in mutant cells (**Figure 6C,D**). Overall, these findings indicate that mutation of Med14S983 triggers feedback mechanisms that compensate for decreased mediator activity and elongation-dependent gene transcription downstream of cAMP signaling.

**Figure 6:**
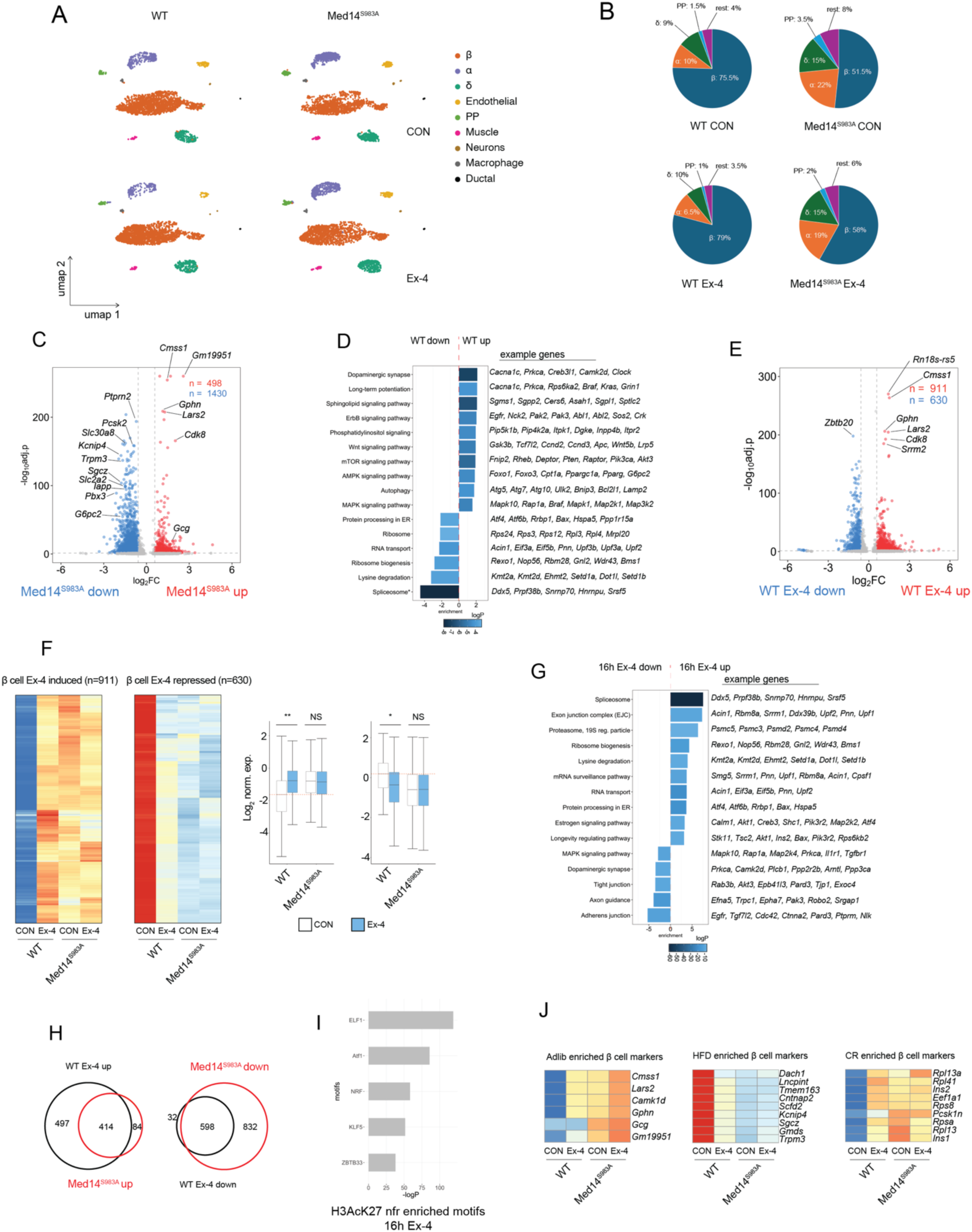
Med14S983 controls beta cell plasticity and gene activation by Ex-4 in primary mouse islet tissue. (A) Uniform manifold approximation and projection (UMAP) representation of WT and Med14 S983A mutant pancreatic islets untreated (top) or treated for 16 hours with 10 nM Ex-4 (bottom). 2,500 nuclei are shown per sample. Cell Type annotations are indicated. (B) Relative cell type composition per condition. (C) Volcano plot comparing pseudobulk gene expression in WT and Med14 S983A mutant β cells. Relevant differentially expressed genes (FC >= 1.5; P Adj. <= 0.05) are highlighted. (D) KEGG pathway enrichment of differential genes in WT and Med14 S983A mutant β cells. Example genes for each pathway are shown on the right. (E) Volcano plot comparing pseudobulk gene expression by by Ex-4 (10 nM) in β cells (16 hours). Relevant differentially expressed genes (FC >= 1.5; P Adj. <= 0.05) are highlighted. (F) Heatmap of genes induced (left) and repressed (middle) by Ex-4. Relative expression in WT and Med14 S983A mutant beta cells is shown. Boxplots showing Log2 expression values of signal shown in the heatmaps. * p < 10^−27^, ** p < 10^−57^ by two-tailed t-test (G) KEGG pathway enrichment of genes regulated by Ex-4 (10 nM) in WT β cells (16 hours). Example genes for each pathway are shown on the right. (H) Overlap between genes significantly induced in Med14 S983A mutant cells (WT CON/Med14 S983A CON: FC <= 1.5; P Adj. <= 0.05) and genes significantly induced by Ex-4 in WT cells (16 hours) (WT Ex-4/WT CON: FC >= 1.5; P Adj. <= 0.05) (left). Overlap between genes significantly repressed in Med14 S983A mutant cells (WT CON/Med14 S983A CON: FC >= 1.5; P Adj. <= 0.05) and genes significantly repressed by Ex-4 in WT cells (16 hours) (WT Ex-4/WT CON: FC <= 1.5; P Adj. <= 0.05) (right). (I) Enrichment of TF binding motifs in nucleosome-free regions (nfr) of H3K27Ac decorated enhancers annotated to beta cell genes induced by Ex-4 (16 hours). H3AcK27 ChIP data from whole primary mouse islet tissue (GEO: GSM3604411). (J) Heatmap depicting expression of marker genes of beta cell subclass enriched in adlib fed (left), high fat diet (HFD) fed (middle) and caloric restricted (CR) (right) beta cells [44].

Exposure of cultured islets with Ex-4 for 16 hours resulted in a broad transcriptional response in WT pancreatic beta cells (n=911 Ex-4/CON >=1.5, padj. <=0.05 / n=630 Ex-4/CON <=1.5, padj. <=0.05) (**Figure 6E, Supplementary Table 14**). Consistent with a role of Med14 S983 in Ex-4 mediated transcription, Med14 S983A mutant beta cells were not responsive to Ex-4 treatment (**Figure 6F, Supplementary Table 15**). In agreement with our observations in INS-1 cells, Med14 S983A mutant cells exhibited gene expression profiles corresponding to Ex-4 treated WT cells. Indeed, ∼83% (414/498) of genes induced in Med14 S983A mutant cells (basal) compared to WT are also induced by Ex-4 in WT cells, while ∼41% (598/1430) of genes repressed in Med14 S983A mutant cells compared to WT are also repressed by Ex-4 in WT cells (**Figure 6G,H**). Motif enrichment analysis of NFRs in H3K27Ac active enhancers of mouse islets revealed enrichment of Atf1 motif in enhancers annotated to Ex-4 induced genes (**Figure 6I**), suggesting a role of CREB in Ex-4 induced gene expression in beta cells.

Recent work linked long-term feeding regimens (adlib, caloric restriction (CR), high fat (HF)) to a shift in beta cell transcriptional states in primary islets using single cell RNA-seq[44]. Strikingly, sustained exposure of Ex-4 induced beta cell marker genes that are enriched in CR (e.g. *Rpl13a*, *Rpl41*, Ins1, *Ins2*, *Eef1a*) and adlib (e.g. *Cmss1*, *Lars2*, *Camk1*) mice and depleted marker genes enriched in and HF (e.g. *Dach1*, *Lncpint*, *Tmem163*, *Cntnap2*) mice in a Med14 dependent manner (**Figure 6C,E,J**). These findings indicate that sustained Ex-4 shifts beta cells to an active metabolic state that favors beta cell survival[44], while confirming a role of Med14 S983 in regulating insulin gene induction by Ex-4 in primary beta cells (**Figure 6J**).

Finally, we tested the role of Med14 S983 in transcriptional response to FSK (16 hours) in primary islets. The role of Med14S983 in FSK induced gene regulation appears more limited in primary beta cells (**Figure S6B, Supplementary Table 16,17**). However, genes induced by 16 hour exposure to FSK in a Med14 S983 dependent manner drive relevant processes like vesicle trafficking (*Astn2, Hid1, Rab20*), actin rearrangement (*Bcar1, Peak1, Ctnnal1*), calcium mobilization (*Ryr3*), protein folding in the ER (*Pdia6*), growth (*Shc4*) and PKA activity (*Crybg3)* in beta, but not alpha, delta or PP cells (**Figure S6C**). Furthermore, a number of genes in this set (*Peak1*, *Astn2*, *Bcar1, Ctnnal1, Ppfibp2*) is associated with significant diabetes-related risk loci (**Figure S6D**)[45]. ChIP-seq analysis in primary mouse islets confirmed CREB binding in active enhancers in Med14 S983 dependent diabetes-associated loci *Bcar1/Ctrb1* and *Peak1/Hmg20a* (**Figure S6E**). Previous work showed that the common SNP rs7202877 in the *Bcar1/Ctrb1* locus significantly associates with decreased GLP-1 induced insulin secretion[46]. Further inspection of the *Bcar1/Ctrb1* locus in INS-1 cells revealed a CREB bound inducible enhancer that was inhibited in Med14 S983A mutant cells, directly confirming the role of Med14 S983 in promoting FSK-induced gene induction in the *Bcar1/Ctrb1* locus (**Figure S6F,S6G**). These observations are consistent with the identification of rs7202877 as an expression quantitative trait locus (eQTL) controlling expression of *Ctrb1,2* in humans[46]. Given the abundance of disease-associated SNPs genomic regions that regulate gene expression in human islets [47–49], these findings link Med14 S983 phosphorylation with sustained Ex-4 treatment in the induction of beta cell-specific transcripts with crucial roles in maintaining glucose homeostasis.

## Discussion

Initially identified as an insulin secretagogue[50, 51], GLP-1 and its stable mimetics were found to amplify the beta cell response to high glucose levels in a sustained fashion. Indeed, GLP-1 induces beta cell adaptation to increased fuel pressure by promoting differentiation, suppressing autophagy, lowering ER stress and rewiring metabolism[52]. These chronic responses were long thought to be transcriptionally driven [4, 5, 12, 13, 53] with CREB being one of the most universally observed mediators of GLP-1 action. Triggering of the GLP-1 receptor in beta cell results in induction of the second messenger cAMP, but how this ubiquitous signaling pathway regulates cellular gene expression in specific cellular contexts is poorly understood.

We found that sustained Ex-4 exposure elicits a broad transcriptional response in INS-1 and primary beta cells that includes induction of genes involved in metabolism, actin remodeling, differentiation and growth factor signaling. We confirmed that this gene response is driven by cAMP, as it is largely mimicked by exposure of the adenylyl cyclase agonist forskolin. In exploring the mechanistic underpinning of the Ex-4 response, we identified the essential mediator scaffolding subunit Med14 as an *in vivo* PKA target, which undergoes Ser983 phosphorylation in response to Ex-4. Upon phosphorylation, Med14 promotes cAMP-induced gene expression, particularly cell-specific delayed-early gene targets. Our genome-wide ChIP and footprinting analyses show that inducible recruitment of CREB and its coactivator CRTC2 to chromatin are reduced in Med14S983A mutant cells despite CREB hyperphosphorylation. H3AcK27 ChIP and ATAC-seq experiments indicate that Med14 phosphorylation plays a crucial role in activation of cAMP-responsive enhancers, allowing subsequent binding of CREB/CRTC2.

Beta cells respond to chronic hyperglycemia by promoting glycolytic and lipogenic pathways. Our data show that Ex-4 promotes lipogenesis in INS-1 cells at least in part by regulating SREBP activity. Med14 S983A mutation impairs regulation of SREBP target gene expression both by Ex-4 and cholesterol, showing that a single residue on mediator controls lipid homeostasis through both nutrient and hormonal inputs. Given the central role of cAMP-PKA signaling in lipid metabolism, future studies should illuminate the role of Med14 phosphorylation on lipogenesis in other metabolic tissues.

Our single nucleus RNA-seq data in primary mouse islets confirmed a broad transcriptional response after prolonged Ex-4 exposure in primary beta cells. In particular, we observed a shift in endocrine cell composition of Med14 S983A knock-in mutant primary islets, with beta cells being depleted while alpha cells are enriched in Med14 S983A compared to WT islets, resembling endocrine cell ratios in islets of Type 2 diabetic mice. Med14 S983A mutant beta cells are also depleted in marker genes recently identified in a beta cell subclass that expands upon high fat diet feeding, suggesting Med14 phosphorylation may play a role in the adaptive response to nutrient status.

Med14 S983 is contained within a 200 amino acid Intrinsically disordered Region IDR. IDRs have been shown to promote the formation of transcriptionally hyperactive condensates containing RNA PolII and coactivators that selectively drive expression of cell-restricted genes by activating distal enhancers. Mounting evidence supports a role for amino acid and RNA charge balance in promoting compartmentalization and transcriptional activity [54–56]. Specifically, localized enrichment of charged residues in Med1 IDRs was found critical for partitioning of activators inside condensates [57]. It is therefore likely that phosphorylation-induced negative charge inside IDRs can alter size, composition and function of molecular condensates in response to hormonal stimuli. The Med14 IDR containing S983 is also enriched in Erk phosphorylation sites that control gene induction by mitogenic stimuli [58]. Future studies should reveal whether Ex-4 and growth factor signaling pathways cooperate on Med14 IDR to establish gene expression programs that promote beta cell growth by altering mediator condensate formation. Altogether our studies unexpectedly reveal that phosphorylation of a single serine residue in the Med14 protein plays a dominant role in the response to GLP-1 agonists and more broadly in metabolic response to hormones.

## Data availability

Raw data can be accessed from GEO under the accession number GSE299399.

## Acknowledgements

This work was supported by NIH grants 5R01 DK083834, R35 CA220538, JDRF Innovative Grant (1-INO-2022-1125-A-N), Paul F. Glenn Foundation for Biology of Ageing Research, The Clayton Foundation for Medical Research, the Leona M. and Harry B. Helmsley Charitable Trust. We thank Maxim Shokhirev for comments on the manuscript and Evan Booker for support with the snRNA-seq experiments.

## Materials and methods

### INS-1 cell culture

INS-1 rat insulinoma cells were grown at 37C with 5% CO2 in RPMI-1640 media (Corning) supplemented with 10% FBS (Gemini), 2 mM glutamine (Mediatech), 1 mM sodium pyruvate (Mediatech), 100 μg/mL penicillin-streptomycin (Mediatech) and 0.05 mM β-mercaptoethanol.

### CRISPR-Cas9 induced targeted mutagenesis and Med19 tagging

Guide RNAs were cloned in BsmBI digested LentiCRISPRv2 (Addgene #52961) as described in Sanjana et al. For Med14 S983A point mutations, HDR templates with 1kb flanking regions were cloned in pUC19 using the Gibson assembly kit. HDR templates contained Ser983Ala mutations, as well as silent mutations for genotyping (HincII site) and for rendering repaired DNA resistant to Cas9 re-digestion. For 3xFLAG/Twin Strep tagging of Med19, 1kb Med19-NTD flanking regions were cloned in AAVS1_Puro_PGK1_3xFLAG_Twin_Strep plasmid (Addgene #68375) as described in Dalvai et al. [59]. 2 ug of both LentiCRISPRv2 and repair template plasmid were transfected in INS-1 cells using Maxcyte ExPERT ATx electroporator with Optimization 2 protocol. After transfection, cells were selected in 1 ug/ml puromycin for three days. Single cells were sorted by FACS in 96 well plates. Genomic DNA from resulting clones was extracted with QuickExtract DNA Extraction Solution (Lucigen) for genotyping.

### Mediator purification

Mediator was purified as described earlier with some modifications. Briefly, 120 15 cm plates of FLAG-Strep-Med19 knocked-in INS-1 cells were washed with PBS, scraped from the plates and centrifuged at 1,500 rpm. Pellets were resuspended in hypotonic buffer (10 mM HEPES (pH7.9), 1.5mM MgCl2, 10mM KCl, 0.5mM DTT, protease inhibitors, phosphatase inhibitors) and incubated in ice for 15 min. Cell suspension was then transferred to a Dounce homogenizer and homogenized with 15-20 strokes with loose pestle. Next, suspension was centrifuged for 20 min at 20,000 rpm at 4C, followed by a gentle wash of the pellet with another 15 ml of hypotonic buffer and additional centrifugation at 5,000 rpm for 8 min. Pellet was resuspended in twice the pellet volume of extraction buffer (20mM HEPES (pH7.9), 1.5mM MgCl2, 0.6M KCl, 0.2 mM EDTA, 0.5 mM DTT, 25% glycerol, protease inhibitors, phosphatase inhibitors) and rotated at 4C for 1 hour to extract nuclear proteins from chromatin. The nuclear extract was centrifuged for 1 hour at 35,000 rpm, 4C. Cleared supernatant was collected and added to M2 anti-FLAG agarose resin twice pre-washed with equilibration buffer (50mM HEPES (pH7.9), 1.5mM MgCl2, 50mM KCl, 0.3M NaCl) and incubated while rotating at 4C overnight. Beads were then washed with 500 ul elution base buffer (50 mM HEPES (pH7.9), 0.1M NaCl, 1.5mM MgCl2, 0.05% Triton X-100) and eluted with elution base buffer containing FLAG peptide.

### Western blotting

Western blotting was performed using a standard procedure. Protein extract was run on polyacrylamide gels at 30 mA per gel and transferred onto Nitrocellulose membranes at 300 mA. Membranes were washed with TBS 0.1% Tween-20 (TBST) and exposed to antibody in TBST with 5% milk overnight at 4C. Membranes were then washed three times in TBST for 15 min. each and incubated for 1 hour with HRP coupled secondary antibody (1/5000 dilution) followed by three more TBST washes. After 10 seconds exposure to SuperSignal™ West Pico PLUS Chemiluminescent Substrate (Thermo Scientific 34578), blots were exposed to imager film and imaged with an X-ray film processor.

### Quantitative PCR

RNA was extracted from cultured cells or tissue with Trizol (Invitrogen 15596026)/chloroform. cDNA was synthesized from 1 μg total RNA using Transcriptor First Strand cDNA Synthesis Kit (Roche 04897030001) according to manufacturer’s instructions. Quantitative PCR was performed using LightCycler 480 SYBR Green I Master mix (Roche 04887352001) in a LightCycler 480 II (Roche).

Guide sequences used for Med14 S983A mutation and N-terminal Med19 tagging (PAM underlined)

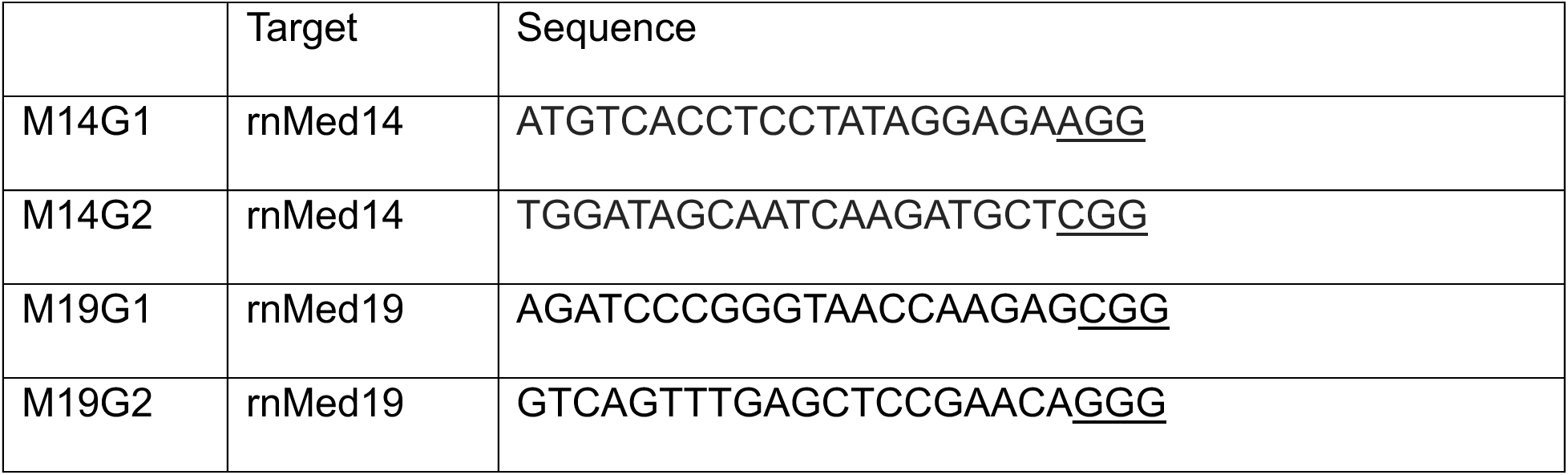

### RNA-seq

Phenol-chloroform extracted total RNA was cleaned up using Direct-zol RNA MiniPrep kit (Zymo Research). 1 ug total RNA was used to build RNA-seq libraries with NEBNext Ultra II Directional RNA Library Prep Kit for Illumina (NEB E7760). 2 ul of 1:100 diluted ERCC Spike-In Mix 1 (Invitrogen 4456740) was added to total RNA. Paired-end sequencing of libraries was performed on the NovaSeq X Plus sequencer (Illumina).

### Chromatin immunoprecipitation (ChIP)

ChIP was performed as described earlier (Heinz, 2010). Briefly, cells were fixed in 0.75% formaldehyde for 10 min and quenched with 125 mM glycine for 5 min. Cells were washed and scraped in ice cold PBS. Cell pellets were resuspended in buffer LB3 (10 mM Tris-HCl pH 8.0, 100 mM NaCl, 1 mM EDTA, 0.5 mM EGTA, 0.1% Na-deoxycholate, 0.5% N-laurylsarcosine, protease inhibitor cocktail, Sigma P9599) and sonicated (Active Motif EpiShear Probe Sonicator). 35 μl protein A agarose beads (Thermo Scientific 20333) were washed twice with PBS + 0.1% BSA and coupled with antibody for 4 hours in PBS + 0.1% BSA at 4°C with rotation. For ChIP, 500 μl extract was added to antibody coupled beads and incubated overnight at 4°C with rotation.

Beads were washed three times in 500 μl wash buffer 1 (20 mM Tris-HCl pH 7.4, 150 mM NaCl, 2 mM EDTA, 0.1% SDS, 1% Triton X-100) and three times in wash buffer 2 (20 mM Tris-HCl pH 7.4, 250 mM LiCl, 1 mM EDTA, 1% Triton X-100, 0.7% Na-deoxycholate). Elution was then performed by incubating beads in 50 μl elution buffer 1 (TE, 1% SDS) for 15 min at 50°C and 50 μl elution buffer 2 (TE, 1% SDS, 300 mM NaCl) for 15 min at 50°C while shaking. Both elutions were combined and incubated with proteinase K and RNase A for 2 hours at 37°C. DNA-protein complexes were de-crosslinked at 65°C while shaking overnight. ChIP DNA was purified using Agencourt AMPure XP beads (Beckman Coulter A63881). 10 ng ChIP DNA was used to build libraries using NEBNext Ultra II DNA library Prep Kit for Illumina (NEB E7645) according to manufacturer’s instructions. ChIPseq reads were aligned to the rn6 reference genome and analyzed for peak detection, read quantification and motif enrichment using HOMER [60]. For detection of nucleosome-free regions in active promoters/enhancers, the following Homer command was used: findPeaks H3K27Ac_tagdirectory -o out -i input_tagdirectory -size 1000 - minDist 2500 -nfr.

### Glycerol gradient ultracentrifugation

Nuclear extracts were loaded on top of a 11 ml 10%-30% linear glycerol gradient in gradient buffer (10 mM HEPES pH 7.4, 100 mM KCl, 1.5 mM MgCl2, 100 uM EGTA, 20 mM NaF, 1 mM DTT, protease inhibitors) and centrifuged in a SW41 rotor at 40,000 rpm for 16 hours at 4C. 500ul fractions were collected from the top.

### Inducible CREB enhancer identification

For identification of inducible CREB enhancers, overlapping CREB bound loci from three independent ChIP-seq experiments (1h FSK exposure) were identified using mergePeaks command in Homer. Next, a raw count table of H3AcK27 ChIP-seq tags over these high confidence CREB bound loci (+/− 2kb) from three independent experiments (basal and 1h FSK exposure) was generated using annotatePeaks.pl. Genomic regions with significantly FSK-induced H3AcK27 were identified using DEseq2 (getDiffExpression.pl) (FC FSK/CON >= 1.5, -log adj. p >= 1).

### ATAC-seq

ATAC-seq was performed as described previously [61], with minor modifications. Briefly, transposition was performed on 50,000 nuclei with Tagment DNA enzyme and buffer for 30 minutes at 37C according to manufacturer’s instructions. DNA was isolated using Qiagen MinElute Reaction Cleanup Kit and amplified with Illumina/Nextera i5 common and i7 index adapters.

ATAC-seq libraries were sequenced on a NovaSeq X (Illumina) (∼2.5E8 reads per library). Reads were trimmed with Trim Galore, aligned to the rn6 reference genome using bwa mem. Mitochondrial, low quality mapped reads and duplicated reads were filtered out with samtools and peaks were called with macs2. Processed fragments were then shifted and sorted into nucleosome-free, mono-, di- and trinucleosome spanning regions with ATACseqQC [62]. Footprinting analysis was performed on short (nucleosome-free) fragments using TOBIAS [63].

### RNAseq data processing

Raw sequencing reads, including ERCC spike-in controls, were mapped to a combined reference genome comprising the rn6 rat genome and ERCC sequences using STAR (v2.5.3a) with default parameters. Following alignment, quality control and gene quantification were conducted using the ‘QoRTs.jar QC’ function from the QoRTs (Quality of RNA-Seq ToolSet) software package (v1.3.6) [64]. This process generated raw expression counts for both ERCC spike-ins and rat genes. Gene expression levels were also quantified in transcripts per kilobase million (TPM) across all exons of RefSeq genes using analyzeRepeats.pl in HOMER (v4.11.1) [60], with the top-expressed isoform used as a proxy for gene expression. Differential gene expression analysis was performed using DESeq2 (v1.24.0) [65] based on raw gene counts, incorporating biological replicates to estimate within-group dispersion. ERCC spike-in abundances were used as quality controls to calculate size factors for normalization in DESeq2. Genes were considered differentially expressed with a false discovery rate (FDR) threshold of <0.05 and an absolute fold-change (FC) >1.5 between experimental conditions.

### Generation of phospho-Med14S983 antisera

Antisera against phospho-Med14S983 were raised in rabbits against a synthetic phosphopeptide corresponding to Med14(974-991) coupled to maleimide activated keyhole limpet hemocyanin per manufacturer’s instructions. The peptide (Cys-DSNQDARRRpSVNEDDNP) was synthesized, HPLC purified and mass spectrometry verified by RS Synthesis (Louisville, KY). The immunogen was prepared by emulsification of Freund’s complete adjuvant-modified Mycobacterium butyricum (EMD Millipore) with an equal volume of phosphate buffered saline (PBS) containing 1.0 mg conjugate/ml for the first two injections. For booster injections, incomplete Freund’s adjuvant was mixed with an equal volume of PBS containing 0.5 mg conjugate/ml. For each immunization, an animal received a total of 1 ml of emulsion in 20 intradermal sites in the lumbar region, 0.5 mg total protein conjugate for the first two injections and 0.25 mg total protein conjugate for all subsequent booster injections. Three individual rabbits were injected every three weeks and were bled one week following booster injections, <10% total blood volume. Rabbits were administered 1–2 mg/kg Acepromazine IM prior to injections of antigen or blood withdrawal. At the termination of study, rabbits were exsanguinated under anesthesia (ketamine 50 mg/kg and aceprozamine 1 mg/kg, IM) and euthanized with an overdose of pentobarbital sodium and phenytoin sodium (1 ml/4.5 kg of body weight IC to effect). After blood was collected the death of animals was confirmed. All animal procedures were conducted by experienced veterinary technicians, under the supervision of Salk Institute veterinarians.

### Affinity purification of phospho-Med14S983 antisera

Antisera were purified using phosphopeptide coupled to SulfoLink coupling resin, 4.6 mg peptide on 2.5 packed bead volume. Coupling was performed by combining peptide and resin in coupling buffer (50 mM Tris, 5 mM EDTA pH 8.5) with 25 mM TCEP and rotating for 1 hour at room temperature followed by alternating washes with 5 column volumes each time of 1 N acetic acid and 50 mM NaHepes, 100 mM NaCl pH 7.5. Next, 20 ml of antiserum was mixed with an equal volume of PBS + 0.02% NaN3, filtered through 5 um and mixed with drained affinity gel by tumbling overnight at 4C. Gel was washed with 5 column volumes of 10 mM Hepes, pH 7.5 and antibody was eluted with 5 column volumes of 1N Acetic acid.

### Oxygen consumption rate (OCR) measurement

INS-1 cells were seeded in XFe 96 microplates at a density of 40,000 cells per well and cultured overnight in RPMI media containing 5 mM glucose. Cells were then washed in serum-free medium without phenol red, sodium bicarbonate, L-glutamine and sodium pyruvate containing 2 mM glucose (Agilent Seahorse XF RPMI Medium, pH 7.4, 103576-100) for 1 hour prior to measurements. Plates were loaded in an XFe 96 Analyzer (Agilent) for OCR measurements. Glucose (20 mM), oligomycin (1 uM) and Antimycin (0.5 uM) + Rotenone (0.5 uM) were injected in the analysis plate at the indicated time points.

### Pancreatic islet isolation

Anesthetized animals were killed by cervical dislocation. Collagenase buffer (HBSS, 2 mM CaCl2, 20 mM HEPES pH 7.4, Collagenase P (0.3 mg/ml)) was injected in the common bile duct and pancreas was digested in a water bath at 37C for 14 min. Digestion was stopped on ice and by addition of 15 ml ice cold HBSS. Pancreases were disrupted by vigorous shaking and filtered through a 500 um mesh. Next, digested tissue was settled by gravity on ice for 10 min. Supernatant was aspirated and tissue washed with HBSS + 10% FBS. Tubes were centrifuged at 220x g for 2 min and dried pellets were resuspended in 10 ml Histopaque. Cell suspension was overlayed with 10 ml HBSS and centrifuged at 850x g for 15 min with brake off. Histopaque containing islets was collected, washed with HBSS + 10% FBS, centrifuged at 220x g for 2 min. Islet pellets were resuspended in 3 ml RPMI + 10% FBS and hand picked three times under a microscope. Islets were cultured overnight and hand picked again before Ex-4 treatment and harvesting.

### Nuclear isolation of pancreatic islets for snRNA-seq

Frozen islets (∼100 per sample) were resuspended in 10 ml of lysis buffer (0.32 M sucrose, 5 mM CaCl2, 3 mM Mg-Acetate, 0.1 mM EDTA pH 8, 10 mM Tris-HCl pH 8, 1 mM DTT, 0.1 % Triton X-100 in DEPC water) supplemented with 10 ul protease inhibitor and 7.5 ul RNase inhibitor (Promega N251B). Suspensions were homogenized on ice in a 15 ml Dounce homogenizer with 15 strokes of the loose (A) and 35 strokes of tight (B) pestle. Nuclei were centrifuged for 5 min at 500g at 4C without brake. Nuclear pellet was resuspended in 400 ul sort buffer (2% BSA, 500 mM EDTA pH 8, 5 ul RNase inhibitor, 1 ug/ml DAPI in dPBS) and filtered through a 30 um CellTric filter into a FACS tube. DAPI-positive nuclei were gated first, followed by exclusion of debris using forward and side scatter pulse area parameters (FSC-A and SSC-A), exclusion of aggregates using pulse width (FSC-W and SSC-W). A BD Influx sorter was used to isolate nuclei, with PBS for sheath fluid (100-μm nozzle was used for cells with sheath pressure set to 20PSI). The nuclei were sorted directly into 1.5mL eppendorf tubes containing 200 ul collection buffer (5% BSA in dPBS, 200U RNase inhibitor) using the “1-drop Pure” sort mode.

### snRNAseq library construction

Isolated nuclei were fixed using Evercode™ Nuclei Fixation v3 (Parse Biosciences ECFN3300) according to manufacturer’s instructions. Combinatorial barcoding of transcriptome on fixed nuclei transcriptome was performed using Evercode™ WT v3 (Parse Biosciences ECWT3300) according to manufacturer’s instructions.

### snRNAseq data processing and analysis

FASTQ files were processed using split-pipe (v1.4.0, Parse Biosciences) to generate cell-by-gene count matrices aligned to the mm39 mouse reference genome. Downstream analysis was conducted using Seurat (v5.3.0)[66]. For each sample, nuclei with a number of counts or detected features in the bottom or top 15th percentile were excluded. Additionally, nuclei with >1% mitochondrial gene content were filtered out. Following quality control, individual samples were merged and normalized using SCTransform [67, 68]. Batch effects were corrected using Harmony [69] prior to clustering.

Cluster-specific marker genes were identified using FindAllMarkers, and the top ten markers from each cluster were submitted to CellKb (https://www.cellkb.com) [70] for annotation. To ensure equal representation across samples, the integrated object was downsampled to 2,500 nuclei per sample before downstream analyses. Differential gene expression between treatment groups and across cell types was performed using FindMarkers (min.pct = 0.1, logfc.threshold = 0). Genes were considered significant with an adjusted p value (BH method) less than 0.05. No fold change cutoff was used. Over-representation analysis was carried out using WebGestaltR (v0.4.6) [71], using the set of all detected genes (prior to filtering) as the reference background.

### Lipid extraction

Lipids were extracted from tissues using the Bligh and Dyer method (Ref). Tissues were homogenized in 1:1 phosphate-buffered saline (PBS):methanol, followed by addition of chloroform to achieve a 1:1:2 PBS:methanol:chloroform solvent ratio (v:v:v). Prior to lipid extraction, the following internal standards were added to chloroform unless otherwise stated: SGDG(13C16-16:0/14:0) for quantification of SGDGs and SGAAGs, and ST(d18:1/17:0) for quantification of sulfatides. The mixtures were shaken vigorously for 30 seconds, vortexed for 15 seconds, and centrifuged at 2,200g for 6 minutes at 4°C. The bottom organic layer was collected and dried under a gentle stream of nitrogen.

### MS sample preparation and analysis

For proteomics samples were precipitated with trichloroacetic acid (TCA, MP Biomedicals, # 196057) overnight at 4 °C or using Methanol-Chloroform. Dried pellets were dissolved in 8 M urea, reduced with 5 mM tris(2-carboxyethyl) phosphine hydrochloride (TCEP, ThermoFisher, #20491), and alkylated with 10 mM iodoacetamide (Sigma, # I1149). Proteins were then digested overnight at 37 °C with trypsin (Promega, # V5111). The reaction was quenched with formic acid at a final concentration of 5% (v/v). Digested samples were analyzed on a Q Exactive Hybrid Quadrupole-Orbitrap Mass Spectrometer.

Samples were injected directly onto a 25 cm, 100 μm ID column packed with BEH 1.7 μm C18 resin (Waters). Samples were separated at a flow rate of 300 nL/min on an EasynLC 1200 (Thermo). Buffer A and B were 0.1% formic acid in water and 90% acetonitrile, respectively. A gradient of 1–10% B over 30 min, an increase to 35% B over 120 min, an increase to 100% B over 20 min and held at 100% B for a 10 min was used for a 180 min total run time.

Peptides were eluted directly from the tip of the column and nanosprayed directly into the mass spectrometer by application of 2.5 kV voltage at the back of the column. The Eclipse was operated in a data dependent mode. Full MS1 scans were collected in the Orbitrap at 120k resolution. The cycle time was set to 3 s, and within this 3 s the most abundant ions per scan were selected for CID MS/MS in the ion trap. Monoisotopic precursor selection was enabled and dynamic exclusion was used with exclusion duration of 60 s.

Protein and peptide identification were done with Integrated Proteomics Pipeline – IP2 (Integrated Proteomics Applications). Tandem mass spectra were extracted from raw files using RawConverter [72] and searched with ProLuCID [73] against Uniprot human database. The search space included all fully-tryptic and half-tryptic peptide candidates. Data was searched with 50 ppm precursor ion tolerance and 600 ppm fragment ion tolerance. Identified proteins were filtered to using DTASelect [74] and utilizing a target-decoy database search strategy to control the false discovery rate to 1% at the protein level [75]. Quantitative analysis of TMT was done with Census [76] filtering reporter ions with 10 ppm mass tolerance and 0.6 isobaric purity filter.

### Global lipidomics analysis

Lipid extracts, normalized by tissue weight, were injected into a Vanquish UHPLC system coupled to a Q-Exactive Plus mass spectrometer (Thermo Fisher Scientific). Lipids were separated on a Waters XBridge BEH C8 column (5 μm, 50 × 4.6 mm) over a 70-minute gradient: 0% B for 5 minutes, 0-20% B in 0.1 minutes, 20-100% B in 50 minutes, held at 100% B for 8 minutes, 100-0% B in 0.1 minutes, and held at 0% B for 6.9 minutes. The flow rate was initially set to 0.1 mL/min, increased to 0.3 mL/min at 5.1 minutes, and further increased to 0.4 mL/min at 63.1 minutes. Data were acquired in positive and negative ionization modes using Xcalibur.

Solvent A consisted of 95:5 water:methanol, and solvent B consisted of 70:25:5 isopropanol:methanol:water. For positive ionization mode, 0.1% formic acid and 5 mM ammonium formate were added to the mobile phases. For negative ionization mode, 0.1% ammonium hydroxide solution (28% NH3 in water) was added. The HESI source parameters were: spray voltage, 3,000 V (positive mode) and 2,250 V (negative mode); capillary temperature, 325°C; sheath gas, 50; auxiliary gas, 10; spare gas, 1; probe temperature, 200°C; S-Lens RF level, 65.

The Q-Exactive Plus mass spectrometer was operated in data-dependent mode with one full MS scan (resolution 70,000, AGC target 1e6, maximum injection time 100 ms, scan range 150-1,500 m/z), followed by ten higher-energy collisional dissociation (HCD) MS/MS scans (resolution 17,500, AGC target 1 × 105, maximum injection time 200 ms, isolation width 1.0 m/z, stepped normalized collision energy of 20, 30, and 40, scan range 200-2,000 m/z). Dynamic exclusion was set to 30 seconds.

Lipids were identified using LipidSearch (Thermo Fisher Scientific) by comparing MS/MS spectra against a built-in database containing over 1.5 million lipid ions and their predicted fragment ions. LipidSearch parameters were: precursor tolerance, 8 ppm; product tolerance, 10 ppm; positive mode adducts, +H, +NH4, +Na, and +H-H2O; negative mode adducts, −H, −2H, and +Cl; m-Score threshold, 5.0. The ID quality filter was set to A and B, requiring annotation of both head group and fatty acyls for confident identification. Maximum lipid intensity was required to be greater than 1 × 105. Additional manual inspection was performed using Skyline56.

### Key resources table

| Reagent or Resource | Source | Identifier |
| --- | --- | --- |
| Antibodies |  |  |
| FLAG (mouse) | Sigma | F1804 |
| Phospho-PKA substrate | Cell Signaling | 9624 |
| Med12 | Cell Signaling | 14360 |
| CDK8 | Cell Signaling | 4106 |
| Med30 | Proteintech | 67038-1 |
| Phospho-CREB (S133) | Cell Signaling | 9198 |
| Fos | Cell Signaling | 32154 |
| JunB | Cell Signaling | 3753 |
| Cpt1a | Abcam | Ab128568 |
| Fasn | Cell Signaling | 3180 |
| SREBP-1a | Santa Cruz | Sc-13551 |
| Ldlr | Abcam | Ab30532 |
| tubulin | EMD Millipore | 05-829 |
| RNAQ Polymerase II CTD repeat phospho S2 | Abcam | Ab5095 |
| Histone H3 (acetyl K27) | Abcam | Ab4729 |
| CRTC2 | In house |  |
| phospho-Med14 (S983) | This study |  |
| Chemicals, peptides |  |  |
| Exenatide (Exendin-4) | Bachem | 4019602 |
| FLAG epitope peptide | Bachem | 4034775 |
| Anti-FLAG agarose | Sigma | A2220 |
| AMPure XP beads | Beckman Coulter | A63881 |
| Collagenase P | Roche | 11213873001 |
| Fetal Bovine lipoprotein deficient serum | Alpha Diagnostic | LDLD47-S |
| Cholesterol | Sigma | C3045 |
| 25-dioxycholesterol | Medchemexpress | HY-113134 |
| Recombinant mouse IL-1β | Gibco | PMC0814 |
| Commercial Kits |  |  |
| NEBNext Ultra II Directional RNA Library Kit for Illumina | New England Biolabs | E7760 |
| NEBNext Ultra II DNA library Prep Kit for Illumina |  | E7645 |
| Micro BCA protein assay kit | Thermo scientific | 23235 |
| Qubit dsDNA HS Assay Kit | Invitrogen | Q32854 |
| Pierce Silver stain kit | Thermo scientific | 24612 |
| Transcriptor First Strand cDNA Synthesis Kit | Roche | 04897030001 |
| LightCycler480 SYBR Green I Master | Roche | 43334520 |
| ERCC RNA Spike-In Mix | Invitrogen | 4456740 |
| Phusion Green High-Fidelity DNA Polymerase | Thermo scientific | F-534L |
| Illumina Tagment DNA Enzyme and Buffer | Illumina | 20034210 |
| NEBuilder HiFi DNA Assembly Master Mix | New England Biolabs | E2621 |
| Evercode WT v3 | Parse Biosciences | ECWT3300 |
| Evercode Nuclei Fixation v3 | Parse Biosciences | ECFN3300 |
| Software and Algorithms |  |  |
| STAR v2.5.3 | Dobin et al. [77] |  |
| MACS2 2.1.2 | Zhang et al. [78] |  |
| ATACseqQC 1.20.2 | Ou et al. [62] |  |
| TOBIAS 0.14.0 | Bentsen et al. [63] |  |
| HOMER 4.11.1 | Heinz et al. [60] |  |
| SAMtools 1.9 | Li et al. [79] |  |
| QoRTs v1.3.6 | Hartley et al. [64] |  |
| DESeq2 1.24.0 | Love et al. [65] |  |

### RT-QPCR oligonucleotides

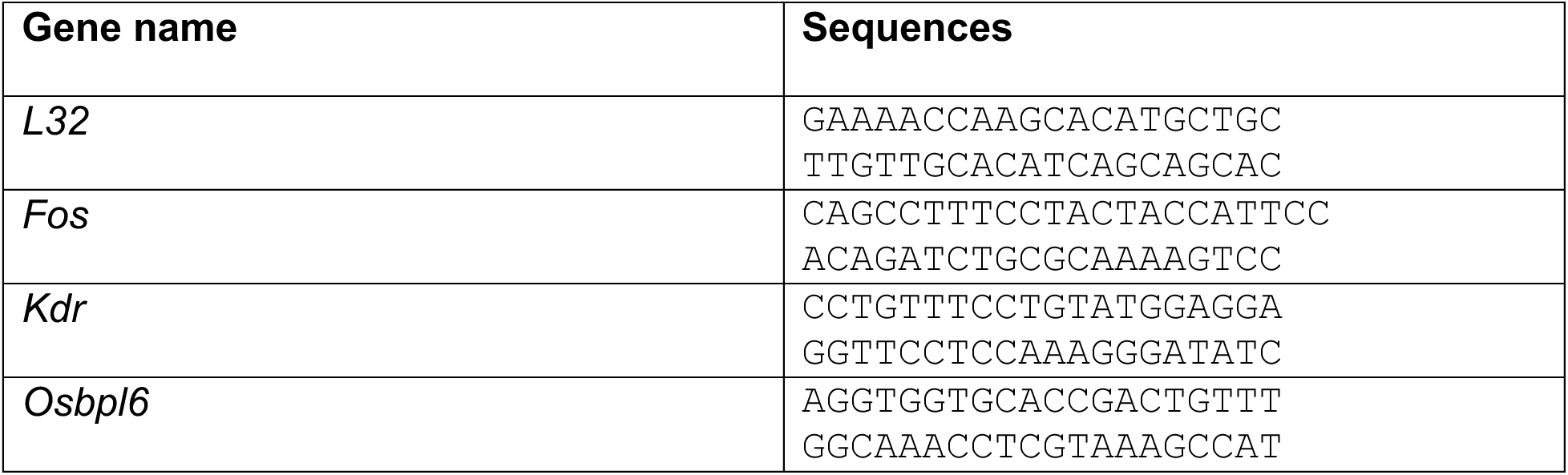
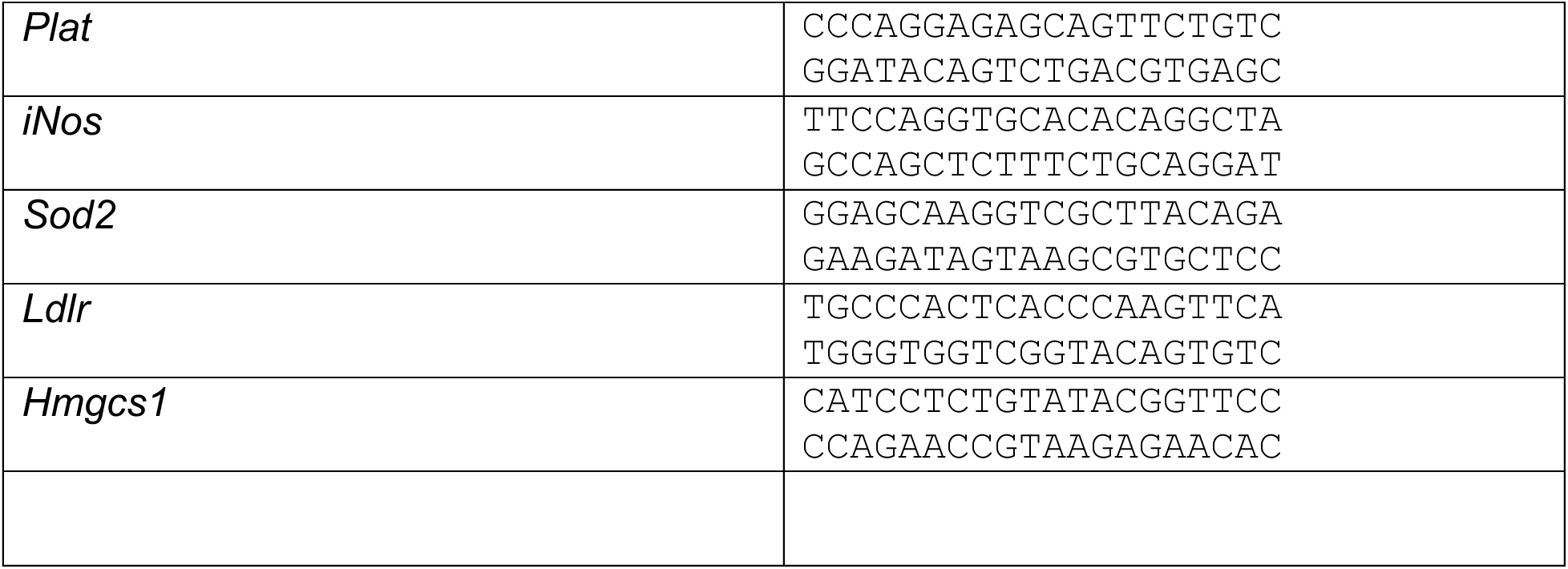

---

**Figure S1:**
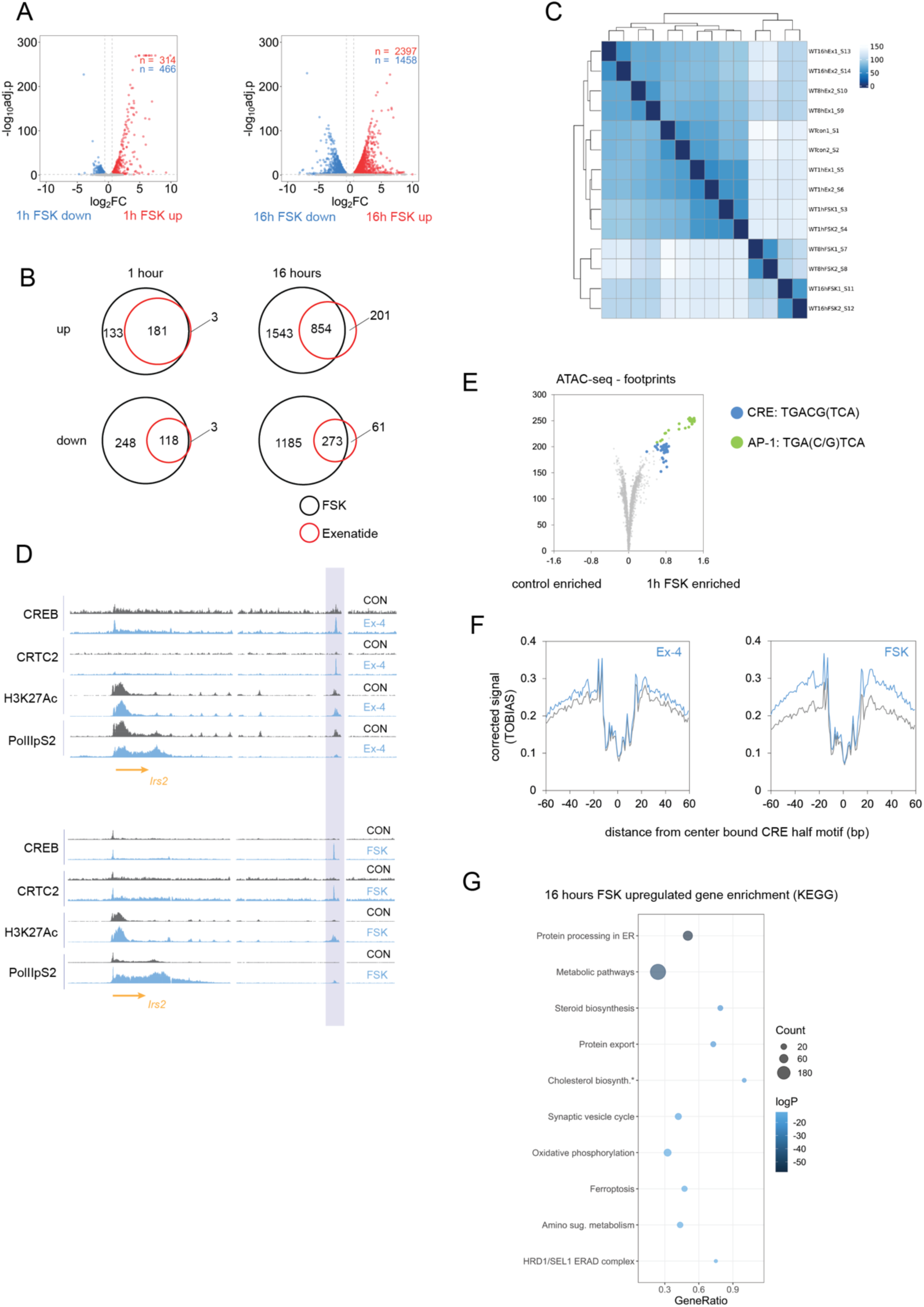
Overlap between Ex-4 and forskolin (FSK) transcriptional responses. (A) Differential gene expression after acute (1h, left) and sustained (16h, right) FSK exposure in INS-1 cells. (B) Overlap of Ex-4 and FSK in induced (top) and repressed (bottom) genes after 1 hour (left) and 16 hour (right) exposure. (C) Sample similarity across all conditions. (D) Example ChIPseq tracks over the *Irs2* locus showing binding of CREB and CRTC induced by Ex-4 (top) and FSK (bottom) over an activated distal enhancer (shaded). Enhancer activity and CTD-phosphorylated RNA polymerase II are shown in H3AcK27 and PolIIpS2 tracks, respectively. (E) Volcano plot depicting change in footprint score over transcription factor binding motifs in accessible chromatin regions (ATAC) after 1 hour FSK treatment. Motifs corresponding to CREB response binding (CRE) and activator protein-1 (AP-1) are highlighted. (F) ATACseq footprints over CRE half (CGTCA) sites after 1 hour Ex-4 (10 nM) (left) and FSK (10 μM) (right) treatment. (G) KEGG pathway enrichment analysis of genes induced after 16 hour FSK (10 μM) exposure.

**Figure S2:**
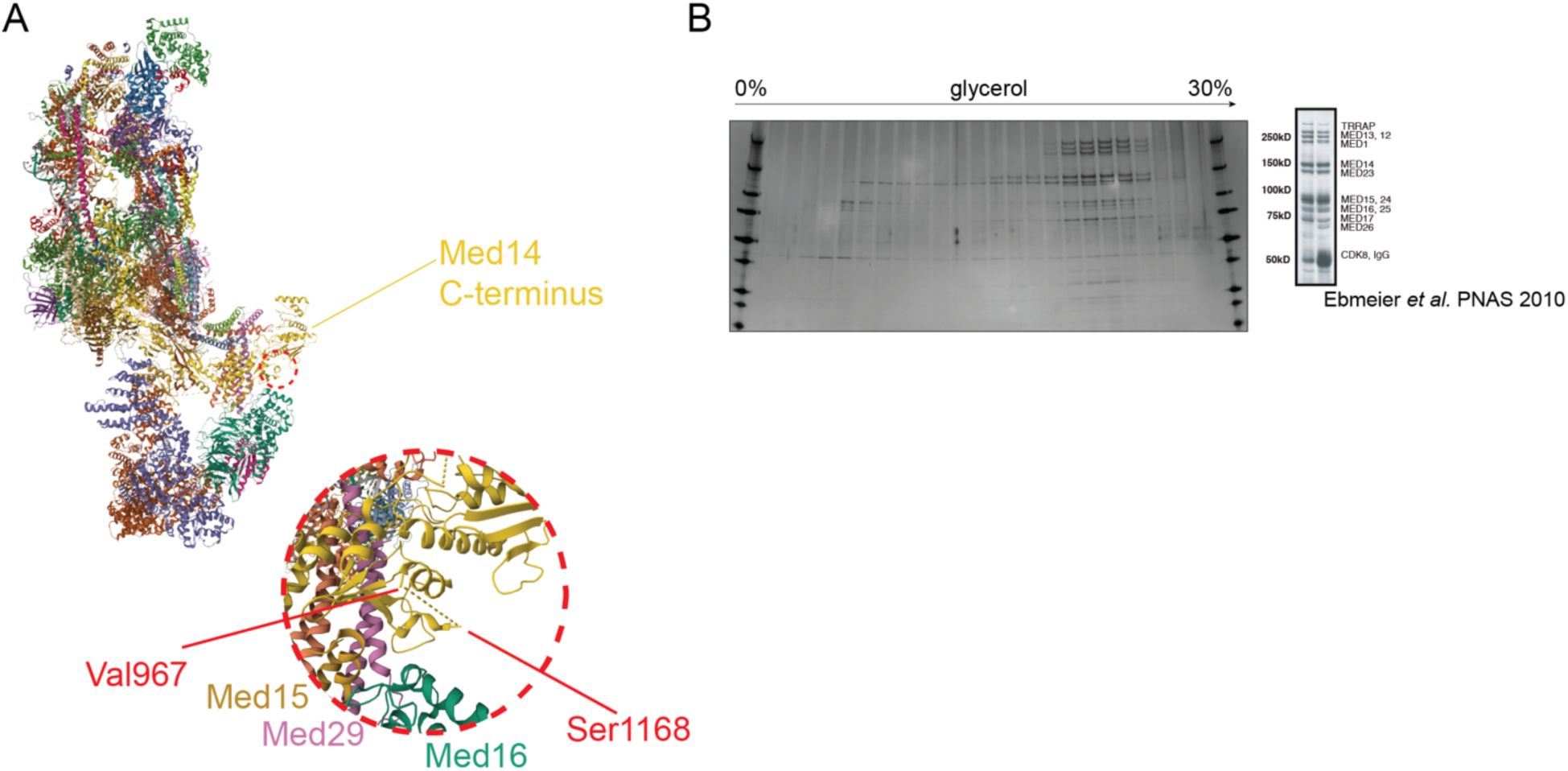
Med14 is a PKA target. (A) Structure of human mediator bound to pre-initiation complex (PDB 7LBM)[16]. Exposed Med14 CTD with unresolved IDR (Med14V967-S1168) is highlighted. (B) Silver stain of purified mediator complex with Med14 S983A loaded on a 10%-30% glycerol gradient. Mediator silver stain form Ebmeier *et al.* [80] shown for reference.

**Figure S3:**
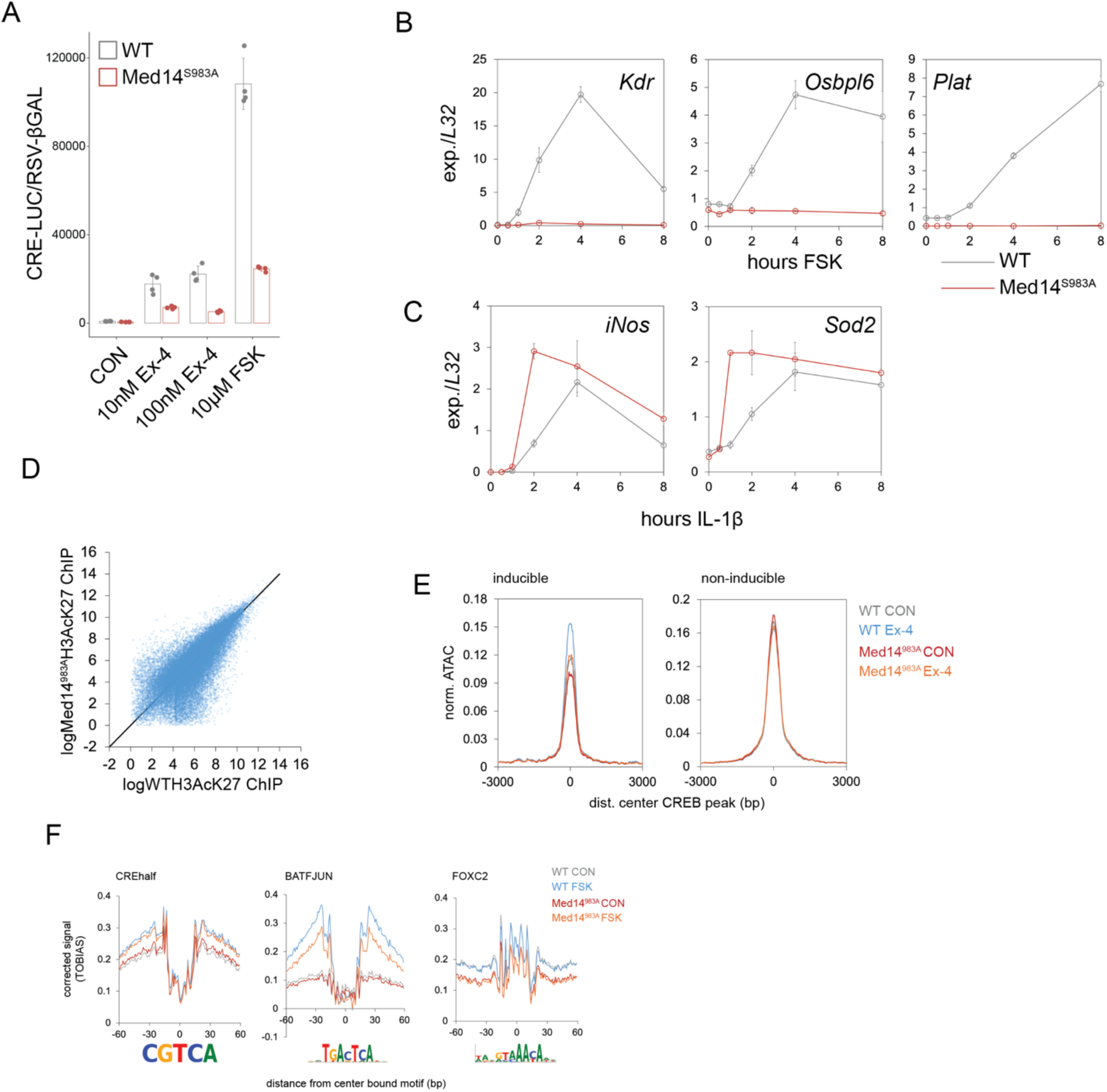
Med14S983 phosphorylation promotes gene induction by activating enhancers. (A) Activation assay of WT and Med14 S983A cells transfected with a CREB-dependent (8xCRE) luciferase reporter. Cells were treated with Ex-4 or forskolin (FSK) as indicated for 6 hours. Signal was normalized by co-transfection of an RSV-βGAL construct. (B) Time course Q-PCR for delayed-early (*Kdr*, *Osbpl6, Plat*) genes over 8 hour FSK (10 μM) treatment in WT and Med14 S983A mutant cells. Error bars show standard deviation. (C) Time course Q-PCR for NF-κB target genes over 8 hour IL-1β (1 nM) treatment in WT and Med14 S983A mutant cells. Error bars show standard deviation. (D) Scatter plot comparing H3AcK27 ChIP-seq reads in enhancers between WT and Med14 S983A mutant cells. (E) Histogram of normalized ATAC-seq reads over inducible (left) and non-inducible (right) enhancers. (F) Footprint profiles over example CRE (CREhalf), AP-1 (BATFJUN) and FOX (FOXC2) motifs in WT and Med14 S983 mutant cells. Cells were exposed to 10 μM FSK for 1 hour.

**Figure S5:**
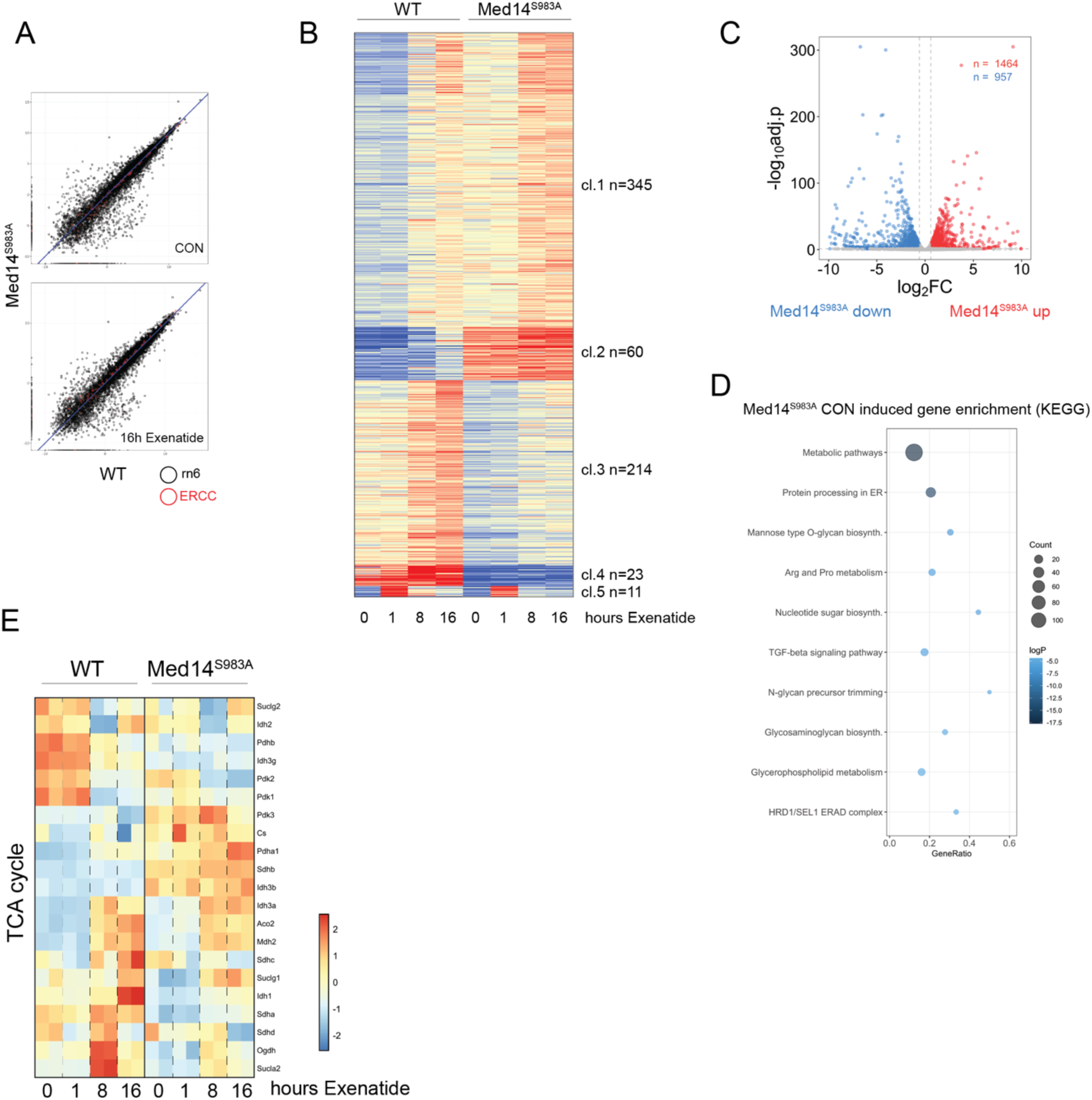
Ex-4 tunes metabolism through Med14. (A) Scatter plot comparing transcript counts between WT and Med14 S983A mutant cells in basal (top) and 16h Ex-4 (bottom) treated conditions. Spiked-in ERCC RNAs are highlighted in red. (B) Clustered heatmap depicting expression of transcripts uniquely induced in WT cells (FC >1.5, Padj. < 0.05) after 16h Ex-4 exposure. Expression across indicated timepoints in WT and Med14 S983A mutant cells is shown. (C) Volcano plot showing differential gene expression in WT and Med14 S983A mutant cells in basal conditions. (D) KEGG enrichment of genes induced in Med14 S983A mutants in basal conditions. (E) Heatmap showing expression of TCA cycle genes in WT and Med14 S983A mutant cells.

**Figure S6:**
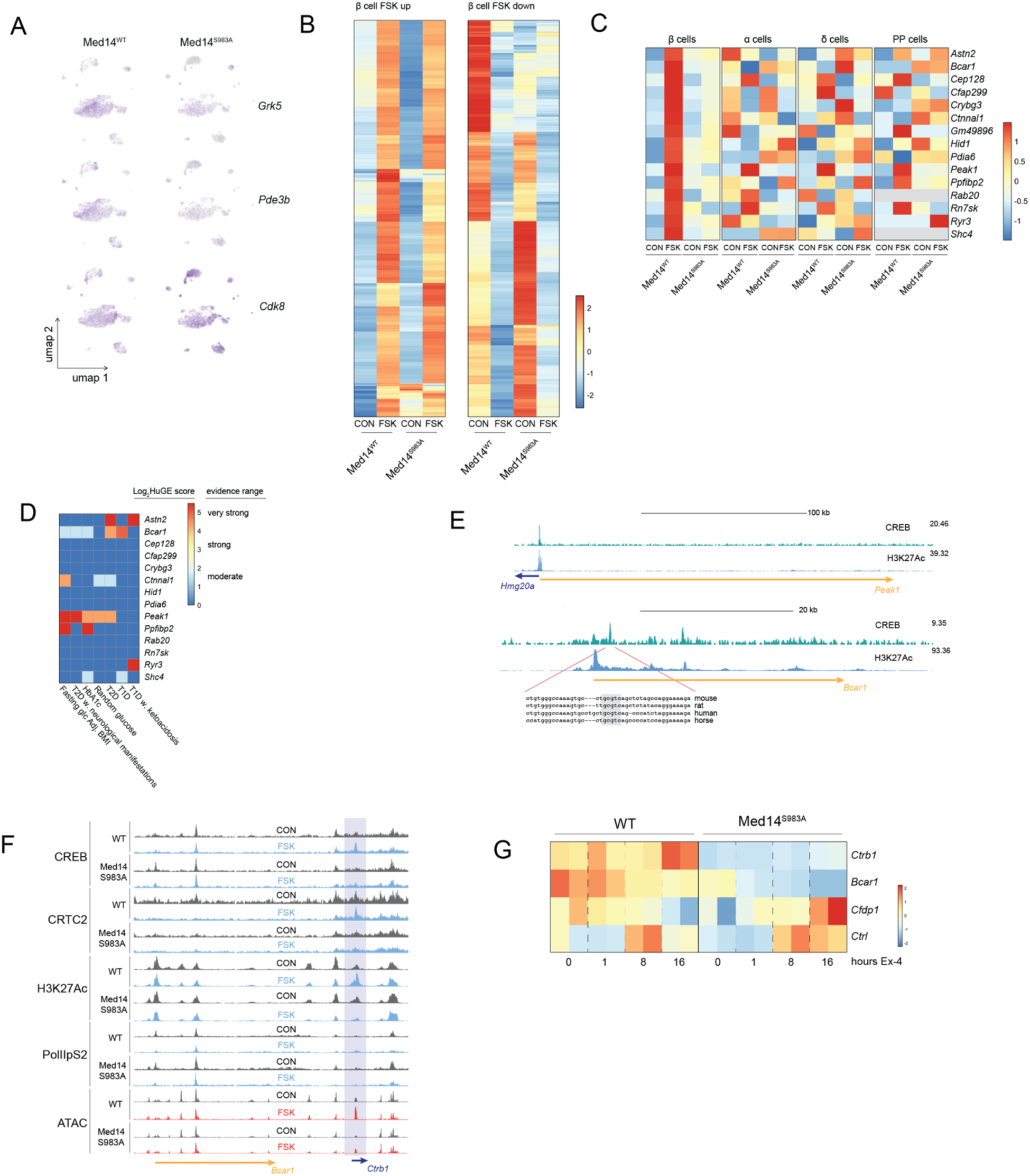
Med14S983 controls beta cell plasticity and gene activation by Ex-4 in primary mouse islet tissue. (A) UMAP plots highlighting expression of negative regulators of cAMP signaling (*Grk5*, *Pde3b*) and mediator kinase subunit *Cdk8*. (B) Gene induction (left) and repression (right) by chronic FSK (16 hours) in WT and Med14 S983A mutant primary beta cells. (C) Heatmap depicting induction of Med14 S983 controlled genes by FSK (16 hours) in WT and Med14 S983A mutants. β, α, δ and PP cells are compared. (D) Heatmap showing Human genetic evidence (HuGE) scores [45] from hugeamp.org for genes shown in (Fig. 6F) across diabetes-related phenotypes. (E) Chromatin occupancy of CREB and H3AcK27 in primary mouse islets across *Hmg20a/Peak1* and *Bcar1* genomic loci in primary mouse islets (GEO: GSM3604407, GSM3604411) [38]. Conserved intronic CREB response element in *Bcar1* intron is shown. (F) ChIP-seq occupancy (top) and ATAC-seq (bottom) tracks over the *Bcar1/Ctrb1* locus in INS-1 cells. cAMP inducible enhancer is highlighted. (G) Expression of genes in the *Bcar1/Ctrb1* locus in INS-1 cells after Ex-4 (10 nM) stimulation.

## References

1. Ashcroft, F.M. and P. Rorsman, Electrophysiology of the pancreatic beta-cell. Prog Biophys Mol Biol, 1989. 54(2): p. 87–143.

2. Prentki, M., F.M. Matschinsky, and S.R. Madiraju, Metabolic signaling in fuel-induced insulin secretion. Cell Metab, 2013. 18(2): p. 162–85.

3. Wortham, M. and M. Sander, Mechanisms of beta-cell functional adaptation to changes in workload. Diabetes Obes Metab, 2016. 18 Suppl 1(Suppl 1): p. 78–86.

4. Van de Velde, S., M.F. Hogan, and M. Montminy, mTOR links incretin signaling to HIF induction in pancreatic beta cells. Proc Natl Acad Sci U S A, 2011. 108(41): p. 16876–82.

5. Carlessi, R., et al., GLP-1 receptor signalling promotes beta-cell glucose metabolism via mTOR-dependent HIF-1alpha activation. Sci Rep, 2017. 7(1): p. 2661.

6. Haythorne, E., et al., Altered glycolysis triggers impaired mitochondrial metabolism and mTORC1 activation in diabetic beta-cells. Nat Commun, 2022. 13(1): p. 6754.

7. Flamez, D., et al., Critical role for cataplerosis via citrate in glucose-regulated insulin release. Diabetes, 2002. 51(7): p. 2018–24.

8. Diraison, F., et al., SREBP1 is required for the induction by glucose of pancreatic beta-cell genes involved in glucose sensing. J Lipid Res, 2008. 49(4): p. 814–22.

9. Wang, H., G. Kouri, and C.B. Wollheim, ER stress and SREBP-1 activation are implicated in beta-cell glucolipotoxicity. J Cell Sci, 2005. 118(Pt 17): p. 3905–15.

10. Drucker, D.J., Biological actions and therapeutic potential of the glucagon-like peptides. Gastroenterology, 2002. 122(2): p. 531–44.

11. Doyle, M.E. and J.M. Egan, Mechanisms of action of glucagon-like peptide 1 in the pancreas. Pharmacol Ther, 2007. 113(3): p. 546–93.

12. Hinke, S.A., K. Hellemans, and F.C. Schuit, Plasticity of the beta cell insulin secretory competence: preparing the pancreatic beta cell for the next meal. J Physiol, 2004. 558(Pt 2): p. 369–80.

13. Jhala, U.S., et al., cAMP promotes pancreatic beta-cell survival via CREB-mediated induction of IRS2. Genes Dev, 2003. 17(13): p. 1575–80.

14. Richter, W.F., et al., The Mediator complex as a master regulator of transcription by RNA polymerase II. Nat Rev Mol Cell Biol, 2022. 23(11): p. 732–749.

15. El Khattabi, L., et al., A Pliable Mediator Acts as a Functional Rather Than an Architectural Bridge between Promoters and Enhancers. Cell, 2019. 178(5): p. 1145–1158 e20.

16. Abdella, R., et al., Structure of the human Mediator-bound transcription preinitiation complex. Science, 2021. 372(6537): p. 52–56.

17. Chen, X., et al., Structures of the human Mediator and Mediator-bound preinitiation complex. Science, 2021. 372(6546).

18. Knuesel, M.T., et al., The human CDK8 subcomplex is a molecular switch that controls Mediator coactivator function. Genes Dev, 2009. 23(4): p. 439–51.

19. Fondell, J.D., H. Ge, and R.G. Roeder, Ligand induction of a transcriptionally active thyroid hormone receptor coactivator complex. Proc Natl Acad Sci U S A, 1996. 93(16): p. 8329–33.

20. Grontved, L., et al., MED14 tethers mediator to the N-terminal domain of peroxisome proliferator-activated receptor gamma and is required for full transcriptional activity and adipogenesis. Mol Cell Biol, 2010. 30(9): p. 2155–69.

21. Yang, F., et al., An ARC/Mediator subunit required for SREBP control of cholesterol and lipid homeostasis. Nature, 2006. 442(7103): p. 700–4.

22. Zhu, Y., et al., Isolation and characterization of PBP, a protein that interacts with peroxisome proliferator-activated receptor. J Biol Chem, 1997. 272(41): p. 25500–6.

23. Yuan, C.X., et al., The TRAP220 component of a thyroid hormone receptor-associated protein (TRAP) coactivator complex interacts directly with nuclear receptors in a ligand-dependent fashion. Proc Natl Acad Sci U S A, 1998. 95(14): p. 7939–44.

24. Ge, K., et al., Transcription coactivator TRAP220 is required for PPAR gamma 2-stimulated adipogenesis. Nature, 2002. 417(6888): p. 563–7.

25. Zamudio, A.V., et al., Mediator Condensates Localize Signaling Factors to Key Cell Identity Genes. Mol Cell, 2019. 76(5): p. 753–766 e6.

26. Sabari, B.R., et al., Coactivator condensation at super-enhancers links phase separation and gene control. Science, 2018. 361(6400).

27. Cho, W.K., et al., Mediator and RNA polymerase II clusters associate in transcription-dependent condensates. Science, 2018. 361(6400): p. 412–415.

28. Jaeger, M.G., et al., Selective Mediator dependence of cell-type-specifying transcription. Nat Genet, 2020. 52(7): p. 719–727.

29. Hnisz, D., et al., A Phase Separation Model for Transcriptional Control. Cell, 2017. 169(1): p. 13–23.

30. Zhang, X., et al., Genome-wide analysis of cAMP-response element binding protein occupancy, phosphorylation, and target gene activation in human tissues. Proc Natl Acad Sci U S A, 2005. 102(12): p. 4459–64.

31. Campbell, J.E. and D.J. Drucker, Pharmacology, physiology, and mechanisms of incretin hormone action. Cell Metab, 2013. 17(6): p. 819–837.

32. Mayr, B. and M. Montminy, Transcriptional regulation by the phosphorylation-dependent factor CREB. Nat Rev Mol Cell Biol, 2001. 2(8): p. 599–609.

33. Glauser, D.A., et al., Transcriptional response of pancreatic beta cells to metabolic stimulation: large scale identification of immediate-early and secondary response genes. BMC Mol Biol, 2007. 8: p. 54.

34. McNeill, S.M., et al., The role of G protein-coupled receptor kinases in GLP-1R beta-arrestin recruitment and internalisation. Biochem Pharmacol, 2024. 222: p. 116119.

35. Lee, S.P., et al., GRK Inhibition Potentiates Glucagon-Like Peptide-1 Action. Front Endocrinol (Lausanne), 2021. 12: p. 652628.

36. Bito, H., K. Deisseroth, and R.W. Tsien, CREB phosphorylation and dephosphorylation: a Ca(2+)- and stimulus duration-dependent switch for hippocampal gene expression. Cell, 1996. 87(7): p. 1203–14.

37. Hagiwara, M., et al., Transcriptional attenuation following cAMP induction requires PP-1-mediated dephosphorylation of CREB. Cell, 1992. 70(1): p. 105–13.

38. Van de Velde, S., et al., CREB Promotes Beta Cell Gene Expression by Targeting Its Coactivators to Tissue-Specific Enhancers. Mol Cell Biol, 2019. 39(17).

39. Sporrij, A., et al., PGE(2) alters chromatin through H2A.Z-variant enhancer nucleosome modification to promote hematopoietic stem cell fate. Proc Natl Acad Sci U S A, 2023. 120(19): p. e2220613120.

40. Iwafuchi-Doi, M., et al., The Pioneer Transcription Factor FoxA Maintains an Accessible Nucleosome Configuration at Enhancers for Tissue-Specific Gene Activation. Mol Cell, 2016. 62(1): p. 79–91.

41. Haythorne, E., et al., Diabetes causes marked inhibition of mitochondrial metabolism in pancreatic beta-cells. Nat Commun, 2019. 10(1): p. 2474.

42. Fujita, Y., et al., Human pancreatic alpha- to beta-cell area ratio increases after type 2 diabetes onset. J Diabetes Investig, 2018. 9(6): p. 1270–1282.

43. Yoon, K.H., et al., Selective beta-cell loss and alpha-cell expansion in patients with type 2 diabetes mellitus in Korea. J Clin Endocrinol Metab, 2003. 88(5): p. 2300–8.

44. Dos Santos, C., et al., Calorie restriction increases insulin sensitivity to promote beta cell homeostasis and longevity in mice. Nat Commun, 2024. 15(1): p. 9063.

45. Dornbos, P., et al., Evaluating human genetic support for hypothesized metabolic disease genes. Cell Metab, 2022. 34(5): p. 661–666.

46. t Hart, L.M., et al., The CTRB1/2 locus affects diabetes susceptibility and treatment via the incretin pathway. Diabetes, 2013. 62(9): p. 3275–81.

47. Parker, S.C., et al., Chromatin stretch enhancer states drive cell-specific gene regulation and harbor human disease risk variants. Proc Natl Acad Sci U S A, 2013. 110(44): p. 17921–6.

48. Su, C., et al., 3D chromatin maps of the human pancreas reveal lineage-specific regulatory architecture of T2D risk. Cell Metab, 2022. 34(9): p. 1394–1409 e4.

49. Fuchsberger, C., et al., The genetic architecture of type 2 diabetes. Nature, 2016. 536(7614): p. 41–47.

50. Drucker, D.J., et al., Glucagon-like peptide I stimulates insulin gene expression and increases cyclic AMP levels in a rat islet cell line. Proc Natl Acad Sci U S A, 1987. 84(10): p. 3434–8.

51. Holst, J.J., et al., Truncated glucagon-like peptide I, an insulin-releasing hormone from the distal gut. FEBS Lett, 1987. 211(2): p. 169–74.

52. Rowlands, J., et al., Pleiotropic Effects of GLP-1 and Analogs on Cell Signaling, Metabolism, and Function. Front Endocrinol (Lausanne), 2018. 9: p. 672.

53. Screaton, R.A., et al., The CREB coactivator TORC2 functions as a calcium- and cAMP-sensitive coincidence detector. Cell, 2004. 119(1): p. 61–74.

54. Patil, A., et al., A disordered region controls cBAF activity via condensation and partner recruitment. Cell, 2023. 186(22): p. 4936–4955 e26.

55. Henninger, J.E., et al., RNA-Mediated Feedback Control of Transcriptional Condensates. Cell, 2021. 184(1): p. 207–225 e24.

56. Sharp, P.A., et al., RNA in formation and regulation of transcriptional condensates. RNA, 2022. 28(1): p. 52–57.

57. Lyons, H., et al., Functional partitioning of transcriptional regulators by patterned charge blocks. Cell, 2023. 186(2): p. 327–345 e28.

58. Galbraith, M.D., et al., ERK phosphorylation of MED14 in promoter complexes during mitogen-induced gene activation by Elk-1. Nucleic Acids Res, 2013. 41(22): p. 10241–53.

59. Dalvai, M., et al., A Scalable Genome-Editing-Based Approach for Mapping Multiprotein Complexes in Human Cells. Cell Rep, 2015. 13(3): p. 621–633.

60. Heinz, S., et al., Simple combinations of lineage-determining transcription factors prime cis-regulatory elements required for macrophage and B cell identities. Mol Cell, 2010. 38(4): p. 576–89.

61. Buenrostro, J.D., et al., Transposition of native chromatin for fast and sensitive epigenomic profiling of open chromatin, DNA-binding proteins and nucleosome position. Nat Methods, 2013. 10(12): p. 1213–8.

62. Ou, J., et al., ATACseqQC: a Bioconductor package for post-alignment quality assessment of ATAC-seq data. BMC Genomics, 2018. 19(1): p. 169.

63. Bentsen, M., et al., ATAC-seq footprinting unravels kinetics of transcription factor binding during zygotic genome activation. Nat Commun, 2020. 11(1): p. 4267.

64. Hartley, S.W. and J.C. Mullikin, QoRTs: a comprehensive toolset for quality control and data processing of RNA-Seq experiments. BMC Bioinformatics, 2015. 16(1): p. 224.

65. Love, M.I., W. Huber, and S. Anders, Moderated estimation of fold change and dispersion for RNA-seq data with DESeq2. Genome Biol, 2014. 15(12): p. 550.

66. Hao, Y., et al., Dictionary learning for integrative, multimodal and scalable single-cell analysis. Nat Biotechnol, 2024. 42(2): p. 293–304.

67. Lause, J., P. Berens, and D. Kobak, Analytic Pearson residuals for normalization of single-cell RNA-seq UMI data. Genome Biol, 2021. 22(1): p. 258.

68. Choudhary, S. and R. Satija, Comparison and evaluation of statistical error models for scRNA-seq. Genome Biol, 2022. 23(1): p. 27.

69. Korsunsky, I., et al., Fast, sensitive and accurate integration of single-cell data with Harmony. Nat Methods, 2019. 16(12): p. 1289–1296.

70. Patil, A.P., A., CellKb Immune: a manually curated database of mammalian hematopoietic marker gene sets for rapid cell type identification. bioRxiv, 2020.

71. Elizarraras, J.M., et al., WebGestalt 2024: faster gene set analysis and new support for metabolomics and multi-omics. Nucleic Acids Res, 2024. 52(W1): p. W415–W421.

72. He, L., et al., Extracting Accurate Precursor Information for Tandem Mass Spectra by RawConverter. Anal Chem, 2015. 87(22): p. 11361–7.

73. Xu, T., et al., ProLuCID: An improved SEQUEST-like algorithm with enhanced sensitivity and specificity. J Proteomics, 2015. 129: p. 16–24.

74. Tabb, D.L., W.H. McDonald, and J.R. Yates, 3rd, DTASelect and Contrast: tools for assembling and comparing protein identifications from shotgun proteomics. J Proteome Res, 2002. 1(1): p. 21–6.

75. Peng, J., et al., Evaluation of multidimensional chromatography coupled with tandem mass spectrometry (LC/LC-MS/MS) for large-scale protein analysis: the yeast proteome. J Proteome Res, 2003. 2(1): p. 43–50.

76. Park, S.K., et al., Census 2: isobaric labeling data analysis. Bioinformatics, 2014. 30(15): p. 2208–9.

77. Dobin, A., et al., STAR: ultrafast universal RNA-seq aligner. Bioinformatics, 2013. 29(1): p. 15–21.

78. Zhang, Y., et al., Model-based analysis of ChIP-Seq (MACS). Genome Biol, 2008. 9(9): p. R137.

79. Li, H., et al., The Sequence Alignment/Map format and SAMtools. Bioinformatics, 2009. 25(16): p. 2078–9.

80. Ebmeier, C.C. and D.J. Taatjes, Activator-Mediator binding regulates Mediator-cofactor interactions. Proc Natl Acad Sci U S A, 2010. 107(25): p. 11283–8.

